# Spatial analysis of cell patterning to aid genetic and phenotypic understanding of grass stomatal density: a case study in maize

**DOI:** 10.1101/2025.05.21.655366

**Authors:** John G. Hodge, Andrew D.B. Leakey

## Abstract

Biological processes involve complex hierarchies where composite traits result from multiple component traits. However, holistically understanding of how sets of component traits interact to underpin genotype-to-phenotype relationships is generally lacking. Stomatal density (SD) is a tractable model system for exploring how high-throughput phenotyping (HTP) data could be exploited by a new spatial analysis approach to better understand a developmentally and functionally important trait. SD is a composite trait, resulting from various components related to cell identity and size, which are themselves governed by a series of spatio-developmental processes. Data from 180 recombinant inbred lines of maize (*Zea mays* (L.)) were analyzed by a new Stomatal Patterning Phenotype (SPP) to: (1) describe the average spatial probability distribution of the nearest neighboring stomata; (2) derive a core set of component traits related to cell size, cell packing and positional probabilities; (3) build a structural equation model of component traits underlying SD; and (4) identify stomatal patterning quantitative trait loci (QTL). The core set of SPP-derived traits explained 74% of the variation in SD. Analyzing SPP component traits allowed some loci previously identified as generic SD QTL to be recognized as specific to lateral versus longitudinal elements of stomatal patterning. Therefore, this study highlights how novel insights can be gained by decomposing a composite trait (e.g. SD) into a set of component traits that were present in HTP data but not previously exploited.

## Introduction

Organisms are structured systems possessing traits that interact within and across hierarchical levels of biological organization including genetic, molecular, biochemical, developmental and physiological processes (Trewavas, 2006). Traits vary across these hierarchical scales from more elementary components to higher-order composite traits (Riska, 1989; Stuber *et al*., 1987).

Compared to the components that underlie them, higher-order composite traits are often evolutionarily or economically important, and can be easier to recognize and measure e.g. survival, fecundity, grain yield, days to heading/flowering, biomass production, and nutritional quality (Cao *et al*., 2020; Kovács, 1990; Riska, 1989; Snowder & Fogarty, 2009; Stuber *et al*., 1987; Yang, *et al*., 2014). On the other hand, understanding of composite-component trait relationships can be advantageous for crop improvement by identifying pleiotropic traits that are more heritable and easier to select upon than the composite traits they influence e.g. kernel row number for yield gain in maize (Fernandes & Lipka, 2020; Stuber *et al*., 1987; Sigmon and Vollbrecht 2010; Wang *et al*. 2019). The need to phenotype multiple traits simultaneously increases the effort required to dissect component-composite hierarchies, but high throughput phenotyping (HTP) methods could alleviate this bottleneck (Banerjee *et al*., 2020; Furbank & Tester, 2011; Maimaitijiang *et al*., 2020; Rajurkar *et al*., 2022; Yang *et al*., 2014), creating new opportunities for research and innovation if appropriate analysis frameworks can be developed to use the data.

This study investigates how data from a ML-enabled HTP method can: (1) aid in generating a holistic assessment of a composite-component trait set; and (2) help advance knowledge of genotype-to-phenotype relationships and trait networks. Stomatal density (SD) within the grass species, maize (*Zea mays* (L.)), is an important and tractable study system. SD is important because of the central role of stomata in regulating plant carbon and water relations, disease susceptibility, as well as energy balance (Baresch *et al*., 2019; de Boer *et al*., 2016; Farquhar & Sharkey, 1994; Lawson & McElwain, 2016; Leakey *et al*., 2019;). Engineering reductions in SD provides a promising path to developing C3 and C4 crops that have greater water use efficiency (WUE) (Casson & Gray 2008; Ferguson *et al*., 2024; Lunn *et al*., 2024; Yoo *et al*., 2010). In addition, SD has been studied as a model system for understanding cell fate and patterning, leading to elucidation of a sequence of spatially explicit signaling pathways during leaf development (Ashraf *et al*., 2023; Chen, *et al*., 2009; Doll *et al*, 2023; Facette *et al*., 2015; Guseman, *et al*., 2010; Kamiya *et al*., 2003; Nunes *et al*., 2020; Zeng, *et al*., 2020). At the simplest level, SD can be characterized as the product of three component traits i.e. stomatal index (SI; the number of stomatal complexes relative to the number of other epidermal cells plus stomatal complexes), stomatal complex size and the size of other epidermal cells (Sack & Buckley, 2016). However, dissecting the genetic basis for SD requires investigations of various component processes that determine stomatal patterning. These include where files or mother cells are committed to a prestomatal identity and whether they can retain this canalization across their ontogeny (Zhou *et al*., 2025). Recently, a growing number of machine learning methods have been developed to automatically locate stomata in micrographs and estimate their size (Tan *et al*., 2024), providing rich HTP datasets for investigation in a variety of species that includes grasses (Ferguson *et al*., 2021; Xie *et al*., 2021).

Arabidopsis has been a long-standing model for studying stomatal development (Spiegelhalder & Raissig *et al*., 2021; Torii, 2021). The eudicot broad-leaf morphology commonly features reticulating venation and epidermal cells with “puzzle piece” morphologies, which can combine to produce many different complex patterns of stomata across the epidermis (Conklin, *et al*., 2018; Esau, 1960; Vőfély *et al*., 2018). Consequently, establishing a generalizable method for quantifying the pattern of stomata in dicots has proven difficult (Conklin, *et al*., 2018; Vőfély *et al*., 2018). By contrast, epidermal cells of grasses are arranged in linear files that run longitudinally from leaf base to tip, with a stereotyped arrangement of cells within stomatal files, which are regularly laterally spaced from the midvein to margin (Esau, 1960; Freeling 1992; Vőfély *et al*., 2018; McKown and Bergmann, 2020). This corresponds to a relatively simple grass leaf morphology with parallel veins spaced laterally at regular intervals (Conklin, *et al*., 2018; Esau, 1960, Vőfély *et al*., 2018).

One of the earliest signals in grass stomatal morphogenesis originates from veins within young leaf primordia, radiating laterally to drive the canalization of epidermal stomatal mother cells along the veins flanks (Baresch *et al*., 2019; Facette *et al*., 2015; Nunes *et al*., 2020). Due to their linear venation, these flanking mother cells also typically form into linear files longitudinally along the length of the leaf blade from ligule to tip (Facette *et al*., 2018; Nunes, 2020; Raissig *et al*., 2021; Torii, 2021). Stomatal formation then involves a single asymmetric division of mother cells which produce two daughter cells, the first being an intervening pavement cell and the second a guard mother cell (Ashraf *et al*., 2023; Facette *et al*., 2015; Raissig *et al*., 2016; Stebbins & Shah, 1960). Guard mother cells subsequently recruit subsidiary cells from their lateral pavement neighbors via two additional asymmetric divisions producing the general stomatal complex morphology common to grasses (Esau, 1960; McKown & Bergmann, 2020; Nunes 2020; Spiegelhalder & Raissig *et al*., 2021).

Although the establishment of lateral patterns of veins and their surrounding cells followed by longitudinal processes of division and differentiation of cells has been recognized as the foundation of grass leaf blade development for some time, many elements of the underlying genetics remain poorly understood. The higher vein densities typical of C4 systems like maize could be anticipated to add additional layers of association between not only stomatal file patterning but also internal anatomy (Sedelnikova *et al*., 2018). Applying quantitative genetic techniques can aid in identifying loci controlling specific component processes, but this depends on being able to phenotype the component processes or effective proxies of them. This was not achieved in most prior work, where SD or SI have been the target traits (Baresch *et al*., 2019; de Boer *et al*., 2016; Casson & Gray, 2008; Doheny-Adams *et al*., 2012; Doll *et al*., 2023; Ferguson *et al*. 2021; Xie *et al*. 2021). In a few cases, tessellations have been used to describe the regular arrangements of stomatal, or even trichome patterning phenotypes (Croxdale, 2000, Balkunde *et al*., 2020). Voronoi or other tessellation methods assume all objects within a field of view are known and optimize the edge placement between clusters of nearest neighbor centroids which can have an uncanny resemblance to epithelial tissues (Aurenhammer, 1991; Balkunde *et al*., 2020; Croxdale, 2000; Kaliman *et al*., 2016). This geometric modeling perspective generally does not consider: (1) how these centroids arrived at their present spatial arrangement, or (2) that the biomechanical forces within and between cells may be asymmetric resulting in observed anisotropies that deviate from geometric models (Kaliman *et al*., 2016). Unfortunately, due to the first of these two issues, normalized Voronoi measures typically lack the capacity to capture distinct spatial attributes of lateral and longitudinal positioning. The ideal solution to phenotyping stomatal patterning would be a spatially explicit description of how stomata are arranged relative to each other and the matrix of epidermal cells in which they are embedded, which takes a generalizable form that could be applied across intraspecific and interspecific phenotypic variation. Nearest neighbor analysis (Keller *et al*., 1985) provides a means to quantitatively survey the relative positions of stomata around an average “origin” stomata in the neighborhood of an epidermal micrograph. Analytical approaches such as Structural Equation Modeling (SEM) are ideal for assessing the relationships among a network of component traits while allowing for orthogonal (independent) paths by which they converge on a shared composite trait such as stomatal density (Mi *et al*., 2010; Shipley, 2000). SEM and related approaches have been infrequently used to address similar genetic-phenotypic interactions (Grotzinger, *et al*., 2019; Li *et al*., 2006; Mi *et al*., 2010; Ottaviano & Camussi, 1981; Wright 1920).

Stomatal density has been shown to be polygenic in numerous species (Delgado *et al*., 2019; Ferguson *et al*., 2021; Gailing *et al*., 2008; Xie *et al*., 2021). As a composite trait, it is possible that the multiple small-effect quantitative trait loci (QTL) identified for SD in a given species (including maize, Xie *et al*., 2021) can be dissected into their corresponding component QTL that may exhibit stronger genetic associations and help explain how this SD variation arises at these loci. Additionally, identifying loci which act through distinct mechanisms could maximize the effects of transgressive segregation driving the largest possible phenotypic shifts with the fewest alleles (Wagner, *et al*., 1998) to support breeding for SD phenotypes that improve WUE.

To address the knowledge gaps and opportunities described above, this study aims to:

1. use existing data from a maize B73xMs71 RIL population describing stomatal locations and sizes to derive a Stomatal Patterning Phenotype (hereafter SPP) that quantifies the probability of an ‘average stomatal spatial neighborhood’;
2. extract a series of component traits describing the SPPs for each of the 180 inbred genotypes, and reduce it to a core set of spatial traits using various modeling and multivariate methods;
3. build an SEM to assess the network of relationships among the composite spatial traits and SD;
4. analyze the genetic basis of these component traits alongside SD through QTL mapping to investigate the genetic architecture of separate lateral and longitudinal elements of stomatal patterning.

## Results

### Characterizing the Key Features of the SPP Phenotype

Images of the abaxial epidermis at a central point on the youngest fully expanded leaf (**Fig. 1A,B**) were analyzed to assess stomatal patterning. Rank-ordered nearest neighbors were then identified for all stomata in the field of view while being not immediately adjacent to the edge of the image (see methods). The stomata that are the 1^st^ or 2^nd^ ranked nearest neighbors tend to either lie in the same file of cells as the origin stomata or in a diagonally adjacent position of a nearby cell file (**Fig. 1C,D red and gold arrows**). In contrast, 3^rd^ ranked nearest neighbors are more commonly found in lateral positions several cell files away from the origin stomata (**Fig. 1E, green arrows**). 4^th^ and 5^th^ ranked neighbors followed a similar pattern seen in the 3^rd^ rank although with increasing noise as the search area expands with each rank order (not shown). Normalizing these vectors relative to their origin stomata’s coordinates revealed a diamond-like scatter plot patterning morphology. This scatter plot contained patches where stomata are more or less commonly found, and which vary among inbred lines (**Fig. 2A-D**). Heatmaps that summarize the probability of finding a neighboring stomata at a given location can be generated for single ranks of nearest neighbors (**Supp. Fig. 1**) or averaged across ranks nearest neighbors (**Fig. 2E-H**) to provide an easy method for visualizing and quantifying genetic variation in longitudinal and lateral spacing based on the respective distance of the in-file and side-file hotspots from the origin, plus variation in how rigidly this spacing was enforced (see Supp. Mat.). The use of Manhattan distances likely contributed to the diamond-shaped pattern of cell distribution. However, the observed patterns were significantly different from null Manhattan distance models and, therefore, indicate patterning that is more than just simple geometric packing (**Supp. Mat. & Supp. Fig. 2**). Genotypes with greater heterogeneity in spatial patterning had more diffuse hotspots indicating where stomata were likely to be located (**Fig. 2H versus 2E**). Z019E0052 and Z019E0140 not only represent extreme SPP morphologies but they also have the correspondingly highest (80 per mm^2^) and lowest (33 per mm^2^) SDs, respectively, observed in this population (**Fig. 2G, H**). Laterally and longitudinally flattening SPPs allowed patterning of genotypes to be overlaid providing a clear indication of this relationship to SD which was not evident for size and shape variation in SCL, SCW, or SCA (**Supp Fig. 3**. By contrast to these extreme RIL phenotypes the parental lines, B73 and Ms71, had intermediate SPP morphologies (**Fig. 2E, F**). A variety of traits were extracted from the SPP heatmaps (**Fig. 3; Supp. Fig. 4**).

**Figure 1.**
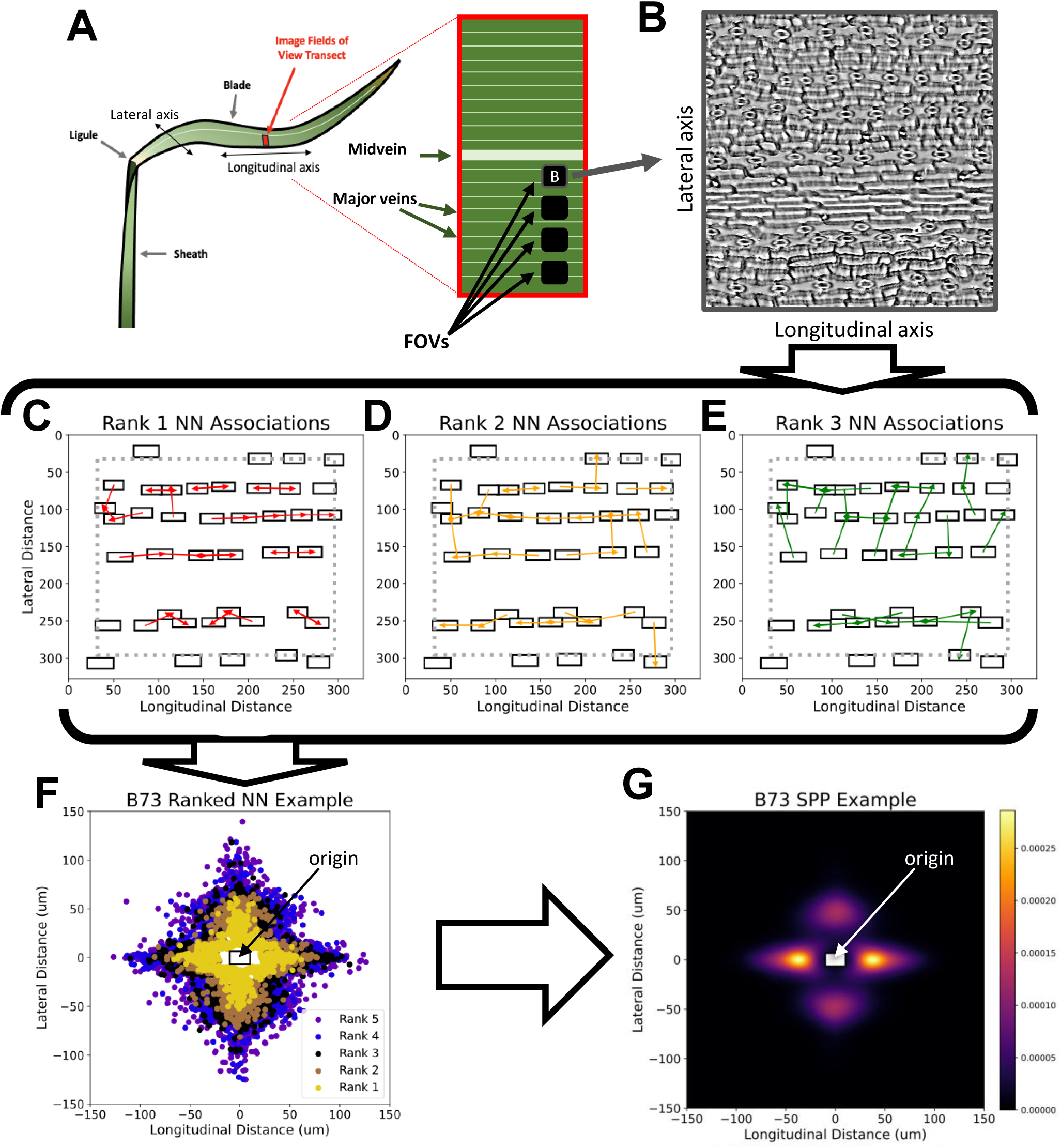
Generalized example of the ranked nearest neighbor phenotyping workflow showing **(A)** the general schema for sample collection from a grass leaf blade and the imaging strategy for fields of view (FOVs) on each sample as well as **(B)** the optical tomographer micrographs generated from this sampling strategy. Ranked nearest neighbor analysis was then performed on the Mask R-CNN annotations of these images with examples of the **(C)** 1^st^ rank, **(D)** 2^nd^ rank, and **(E)** 3^rd^ rank order nearest neighbors for each stomata (white rectangles) denoted as red, gold, and green vectors, respectively. **(F)** These rank-order nearest neighbor vectors were then normalized to their origin stomata’s starting position with an average lengths and widths of these origin stomata being used to generate an average origin stomata (white rectangle). Note that 1st – 5^th^ rank-order nearest neighbors in this scatter plot all have a generally conserved diamond-like distribution, although this pattern diffuses further and further away with sequential rank-orders (i.e., moving from gold, to blue, to purple). **(G)** To generalize these scatterplots into a homologous phenotype which could readily be compared a 2-dimensional kernel density (i.e., a 2D spatial probability) distribution was generated with a set resolution for each tile of the heatmap providing a common Stomatal Patterning Phenotype (SPP) for analysis.

**Figure 2.**
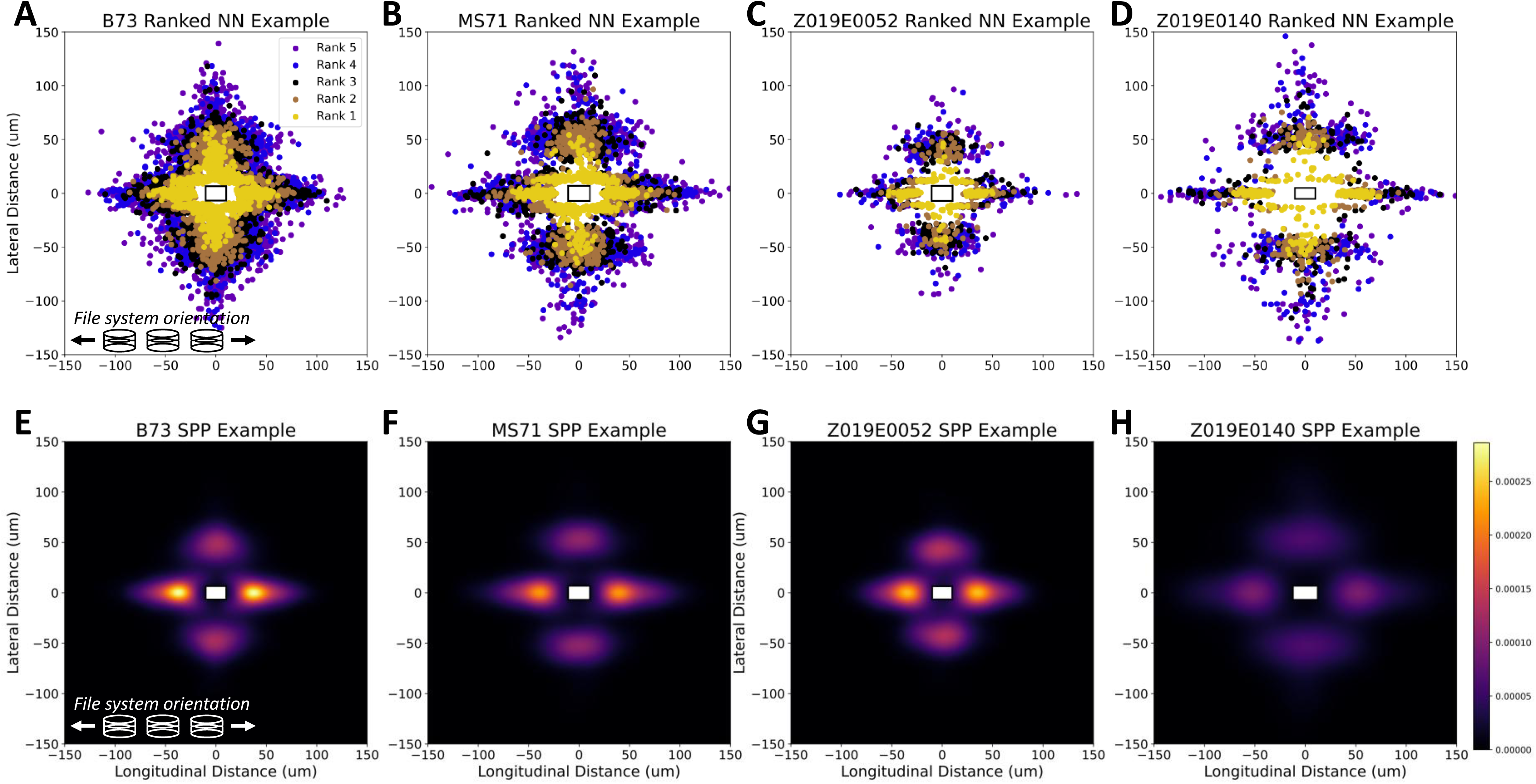
The 1^st^-5^th^ rank ordered nearest neighbors (**A-D**) for the two parental genotypes **(A)** B73 and **(B)** Ms71 in addition two exemplar RILs **(D)** Z019E0052 and **(F)** Z019E0140. Additionally, the corresponding SPP heatmaps which have (**E-H**) that have all been normalized to the same scale which correspond to the two parental genotypes **(E)** B73 and **(F)** Ms71, and the two exemplar RILs **(G)** Z019E0052 and **(H)** Z019E0140. The white rectangle corresponds to the length and width of the average “origin stomata” for each genotype to provide a sense of the relationship between stomatal shape and size and these spatial patterns.

**Figure 3.**
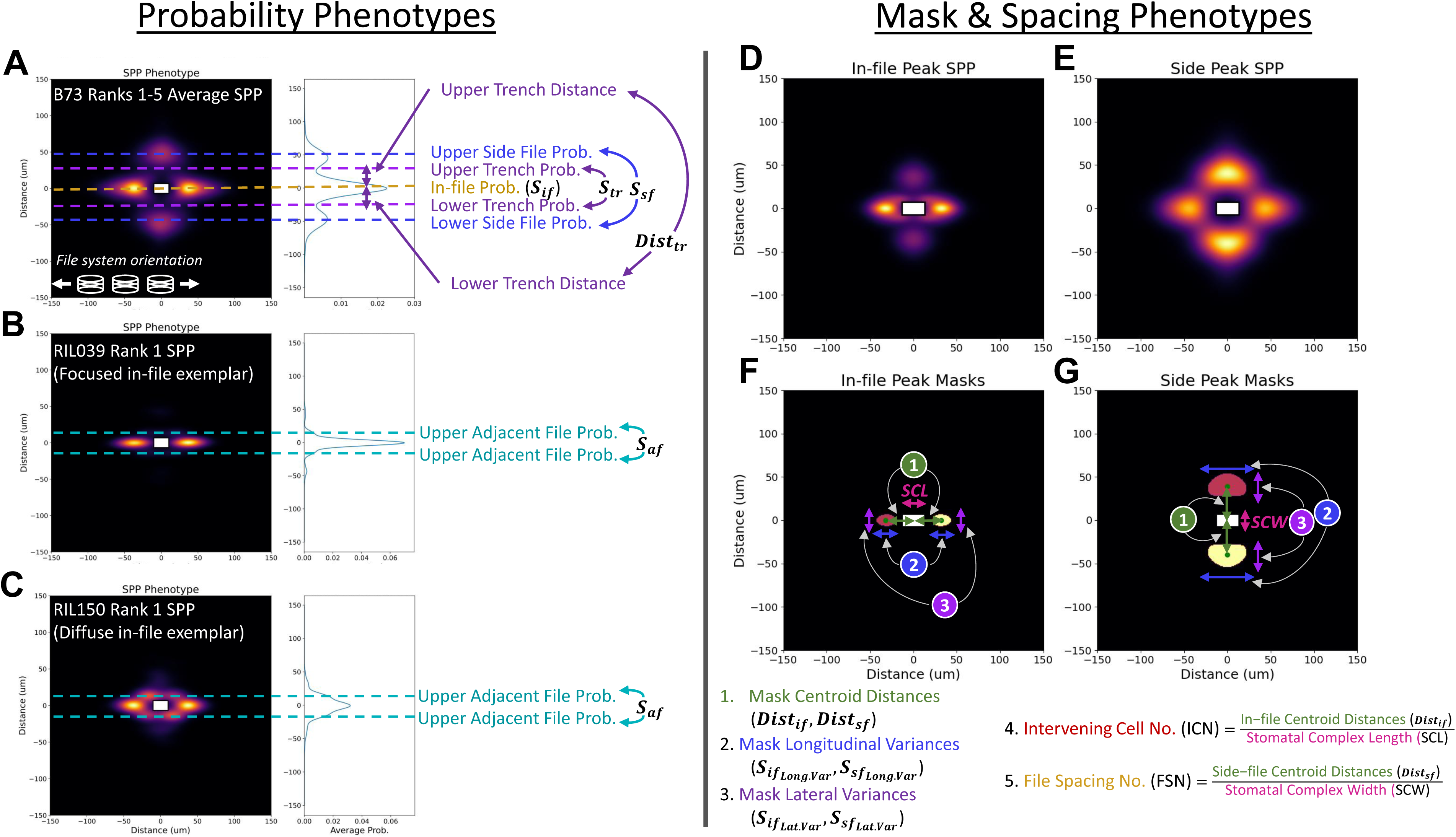
Examples of various SPP traits assessed within this study which can broadly be broken down into two categories **(A-C)** probability phenotypes (left) derived from the longitudinally flattened SPP probabilities measured at discrete vertical positions along the distribution and **(D-G)** mask phenotypes which capture spatial packing information. There are 4 primary probability probabilities that can be assessed, 3 of which are readily retrieved from the 1^st^ - 5^th^ NN averaged SPPs **(A)** which include the in-file probability (dotted gold) falling directly along the origin, the trench probability (dotted purple) which arise as a local minima flanking either side of the this in-file modulus, and finally the side-file probability (dotted blue) which appear as weaker moduli along the outer flanks of each trench. Additionally, the trench distance (black arrows) can be inferred as the distances between the location of the trench probability and the origin. **(B,C)** the fourth probability for adjacent files (dashed grey) is a special case only found among a subset of genotypes in which the first NN for each stomata is typically found immediately at the diagonal **(C)** resulting in a chevron like trimodal pattern to the left and right of the origin which is entirely lacking **(B)** in non-adjacent genotypes. Mask phenotypes (right) are the product of binary masks generated for either the in-file SPP masks **(D,F)** using the 1^st^ and 2^nd^ NN distances or the side-file SPP masks **(E,G)** using the 4^th^ and 5^th^ NN distances. **(1)** Centroid distances (green arrows) can be estimated for both these in-file and side-file masks relative to the origin in addition to the **(2)** longitudinal variance (blue arrows) and **(3)** lateral variance (purple arrows). The in-file and side-file centroid distances can also be normalized relative to the average (pink arrows) stomatal complex length or width to generate an estimated Intervening Cell No. (ICN) or estimated File Spacing No. (FSN), respectively which act as cell packing measures.

### SPP Trait Associations and Identification of Key Sources of Phenotypic Variation

Pairwise correlations, ordination analysis and linear models were used to identify trait-trait associations which could underly key processes in the SPP trait space while eliminating redundant traits from downstream analysis. Within the correlation matrix, specific associations were immediately recognizable such as the significant positive association between *S_if_* and the cell density measures SD or PD (*ρ*_*S*_*if*_,*SD*_ = 0.74, *ρ*_*S*_*if*_,*PD*_ = 0.56 **Fig. 4**). Inversely, the *Dist*_*if*_ exhibited a significant negative relationship with both SD and PD (*ρ*_*Dist*_*if*_,*SD*_ = −0.58, *ρ*_*Dist*_*if*_,*PD*_ = −0.39, **Fig. 4**). When *Dist*_*if*_ is normalized to reflect cell size (i.e., ICN) we see that the magnitude of this negative association is also much weaker (*ρ*_*ICN*,*SD*_ = −0.25, *ρ*_*ICN*,*PD*_ = −0.12, **Fig. 4**). Both cell packing traits ICN and FSN also exhibit modest to weak negative associations with their corresponding normalization factors, SCL and SCW, suggesting that once centroid distances are normalized to cell size, they exhibit a negative relationship with it (*ρ*_*ICN*,*SCL*_ = −0.31, *ρ*_*FSN*,*SCW*_ = −0.2, **Fig. 4**). The probability traits appear to form two distinct groups with *S_if_* and *S_af_* exhibiting similar magnitudes and directions of association while *S_tr_* and *S_sf_* showed a similar relationship amongst themselves (**Fig. 4**). The potential for *S_if_* & *S_af_* and *S_tr_* & *S_sf_* being largely independent of one another was also seen in the minimal correlations between *S_if_* ∼ *S_tr_* or *S_af_* ∼ *S_sf_* (*ρ*_*S*_*if*_,*Str*_ = −0.04, *ρ*_*S_*af*_*,*S_sf_*_ = 0.02 **Fig. 4**). The allometric trait of SLA (a measure of leaf density) was generally found to have the least correspondence to many of the traits except for *S_tr_* where a modest positive correlation was discernible (*ρ*_*S*_*LA*_,*Str*_ = 0.29, **Fig. 4**).

**Figure 4.**
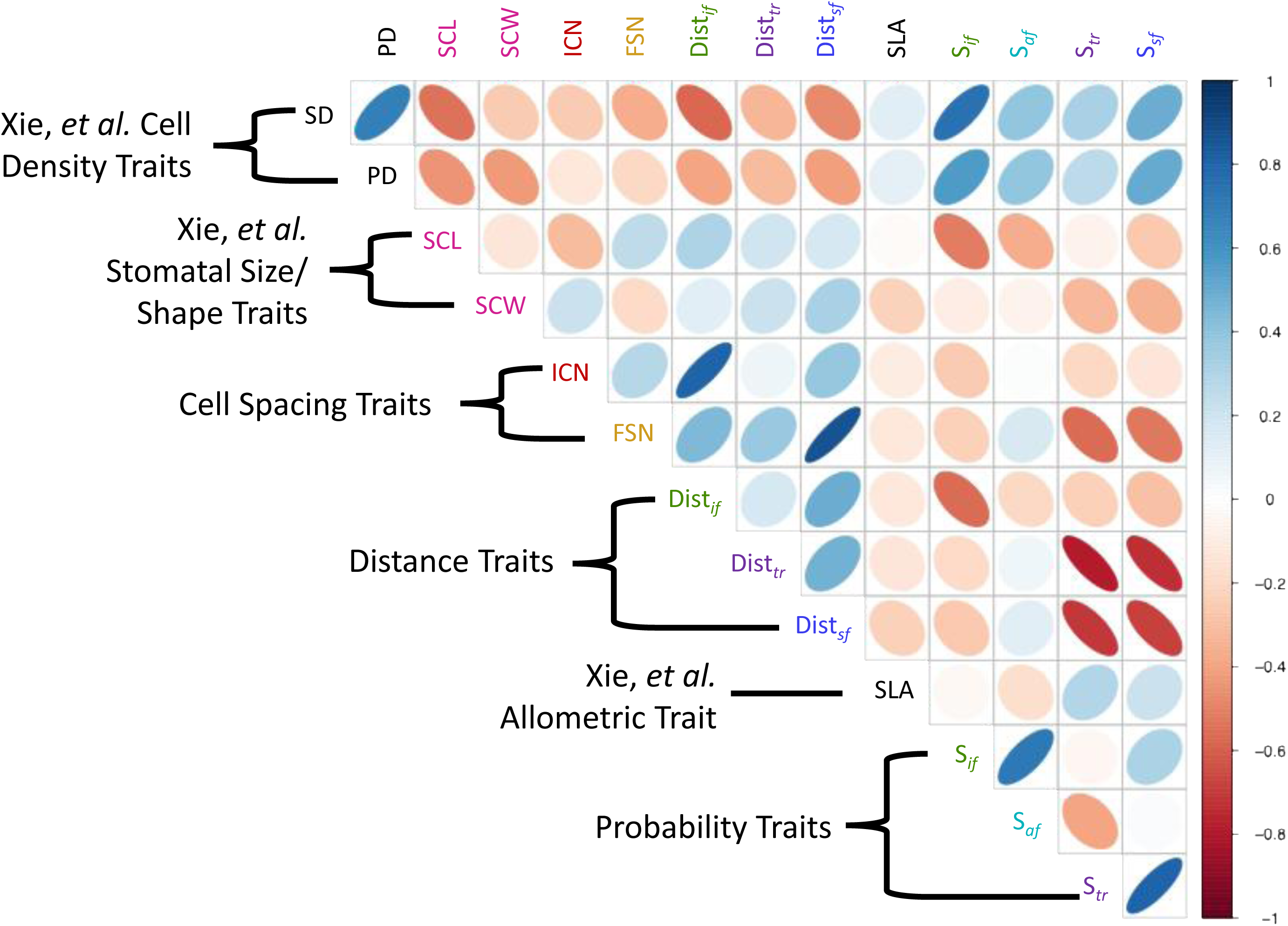
A correlation matrix of the Mask R-CNN and SPP traits. Brackets are provided which denote the class of traits as relating to either cell density, stomatal size/shape, or leaf allometry (these classes all being Xie, *et al*. traits), in addition to cell spacing, distance, or probability traits.

The PCA loadings of the primary probability traits (*S_if_*, *S_tr_*, *S_sf_*), stomatal packing traits (ICN, FSN), and stomatal size traits (SCL, SCW) were generally found to be non-overlapping suggesting that these features were capturing novel information (**Supp. Fig. 5**). By contrast, when traits overlapped in their PCA loadings e.g. the lateral and longitudinal variances for the in-file or side-file probability hotspots and their corresponding centroid distances (*Dist*_*if*_ & *Dist*_*sf*_) the redundant variance traits were dropped. SLA loadings generally tracked with the lateral probability traits *S_tr_* and *S_sf_* (**Supp. Fig. 5**).

Prediction models were then used to refine identification of the minimal trait set that best captured how SD variation arises. Nine models were tested that represented a sequence with progressively greater numbers of traits being used to predict SD (**Supp. Fig. 6A).** The proportion of the variation in SD explained by the models initially increased as more components were added (Models 1-5) but then plateaued (Models 6-9; **Supp. Fig. 6A).** Models 1-3 were limited to inclusion of stomatal shape and size attributes like SCL and SCW (*R^2^*=0.05-0.4, **Supp. Fig. 6B-D**). Models 4-6 then added the SPP cell spacing (ICN, FSN) and probability (*S*_*if*_, *S*_*tr*_, and *S*_*sf*_) traits (*R^2^*=0.57-0.74, **Supp. Fig. 6E-G**). Models 7-9 then added SPP variance traits although this offered negligible improvements (*R^2^*=0.75-0.76, **Supp. Fig. 6H-J**). Although the cell spacing traits ICN and FSN delivered only modest improvements in linear model performance, the PCA revealed they occupy unique regions of the ordination leading to their retention (**Supp. Fig. 5, 6**). Importantly, this model 6 design and its coefficients were able to explain ∼74% of the variation in SD itself (**Supp. Fig. 6G**).

### Structural Equation Modeling of Stomatal Density to Identify Causal Paths of Variation

Structural equation modeling (SEM) described the network of interactions among the core SPP trait set selected above, and ultimately SD. The most likely model (**Fig. 5**) was identified based on AIC values and directed separation tests after several generations of SEM were created to test a variety of hypothetical networks (**Supp. Fig. 7**) assuming SD is a common causal descendant of the other traits. Three hierarchical tiers of causal associations among the traits were identified. Cell spacing traits (ICN and FSN) and stomatal size and shape (SCL and SCW) acted as independent causal drivers. The probability traits *S*_*if*_, *S*_*tr*_, and *S*_*sf*_ act as subordinate traits to cell spacing and size which then feed explanatory variation into SD. The stronger coefficients within this SEM (thicker arrows) reveal anticipated interactions like longitudinally oriented (i.e., SCL→ *S*_*if*_ ←ICN) as well as laterally oriented causal paths (i.e., SCW→ *S*_*tr*_ ←FSN). The trench probability (*S*_*tr*_) appears to have antagonistic paths, being positively associated with *S*_*sf*_ while being strongly, negatively associated with *S*_*if*_. This complex interaction between *S*_*tr*_, *S*_*sf*_, and *S*_*if*_ was not discernible within the pairwise correlations where *S*_*tr*_ and *S*_*if*_had a modest negative correlation despite being the strongest interaction within the SEM model (**Fig. 4**, **5**). Less intuitive patterns also emerged that explain how SD variation arises, with the strongest causal path in this RIL population being FSN→*S*_*tr*_ → *S*_*if*_ →SD, highlighting how lateral and longitudinal elements of stomatal patterning can interact with one another. Ultimately, *S*_*if*_ is the node through which greatest variation in SD arises, as was anticipated from the correlation matrix (**Fig. 5**).

**Figure 5.**
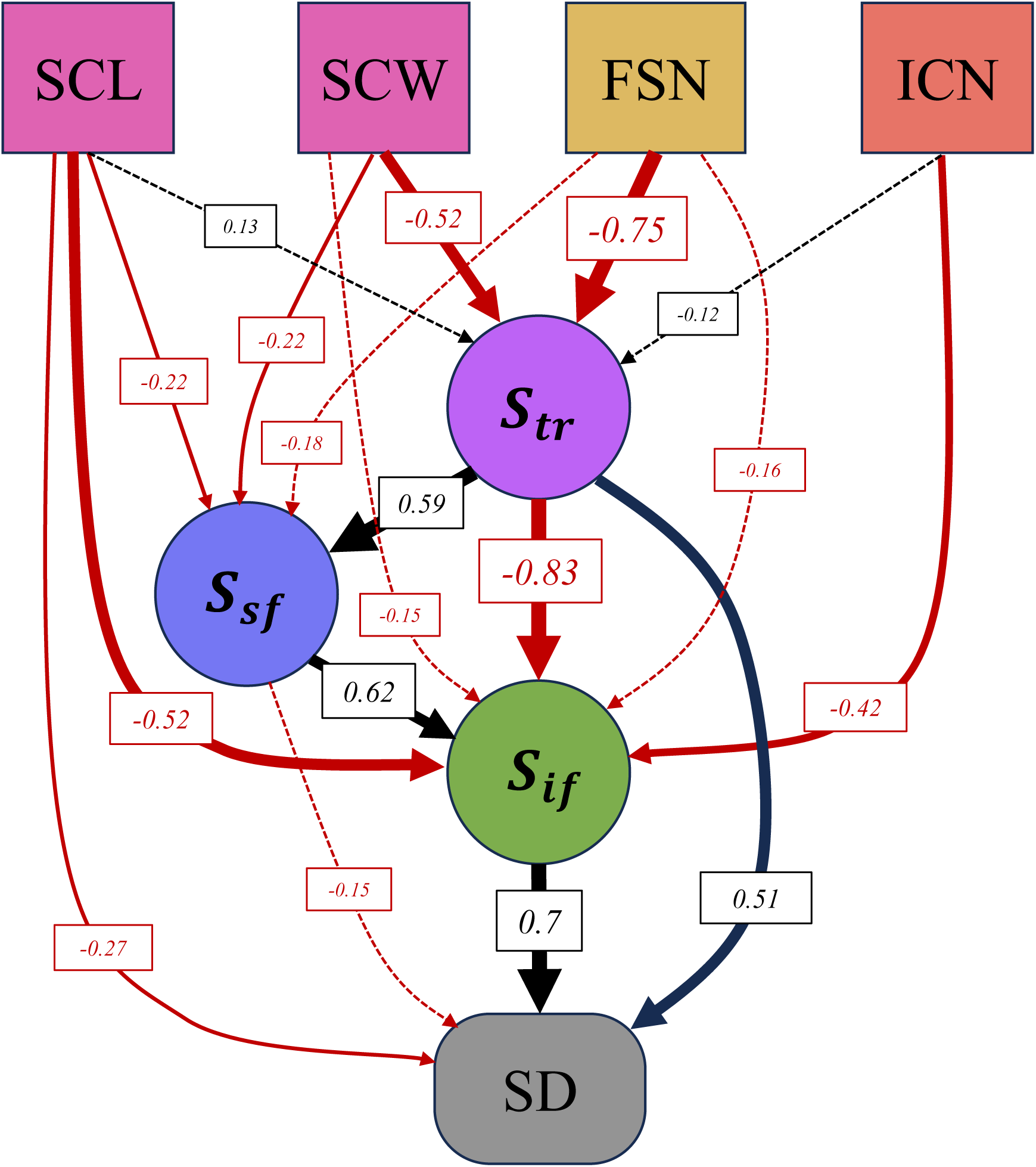
Piecewise Structural Equation Model (SEM) of stomatal patterning and physiological traits shown as a directed acyclic graph of the 2016 field season data in which stomatal shape and size components in addition to the cell file spacing numbers act as common causes for the other traits which ultimately influence SD primarily through acting on either the In-file or Trench probabilities. Additionally, the in-file intervening pavement cell number acts a negative driver of the In-file probability.

### Genetic Mapping of SPP Traits

For traditional measures of stomatal patterning (i.e. SD), QTL mapping recapitulated a previous study with the majority of QTL overlapping exactly (35/41 = 85% congruence with Xie et al. 2021). QTL for SPP traits were generally found to be overlapping with those related to SD and stomatal size and shape and often in hotspots of apparent pleiotropy (**Fig. 6, Supp. Fig 8**).

**Figure 6.**
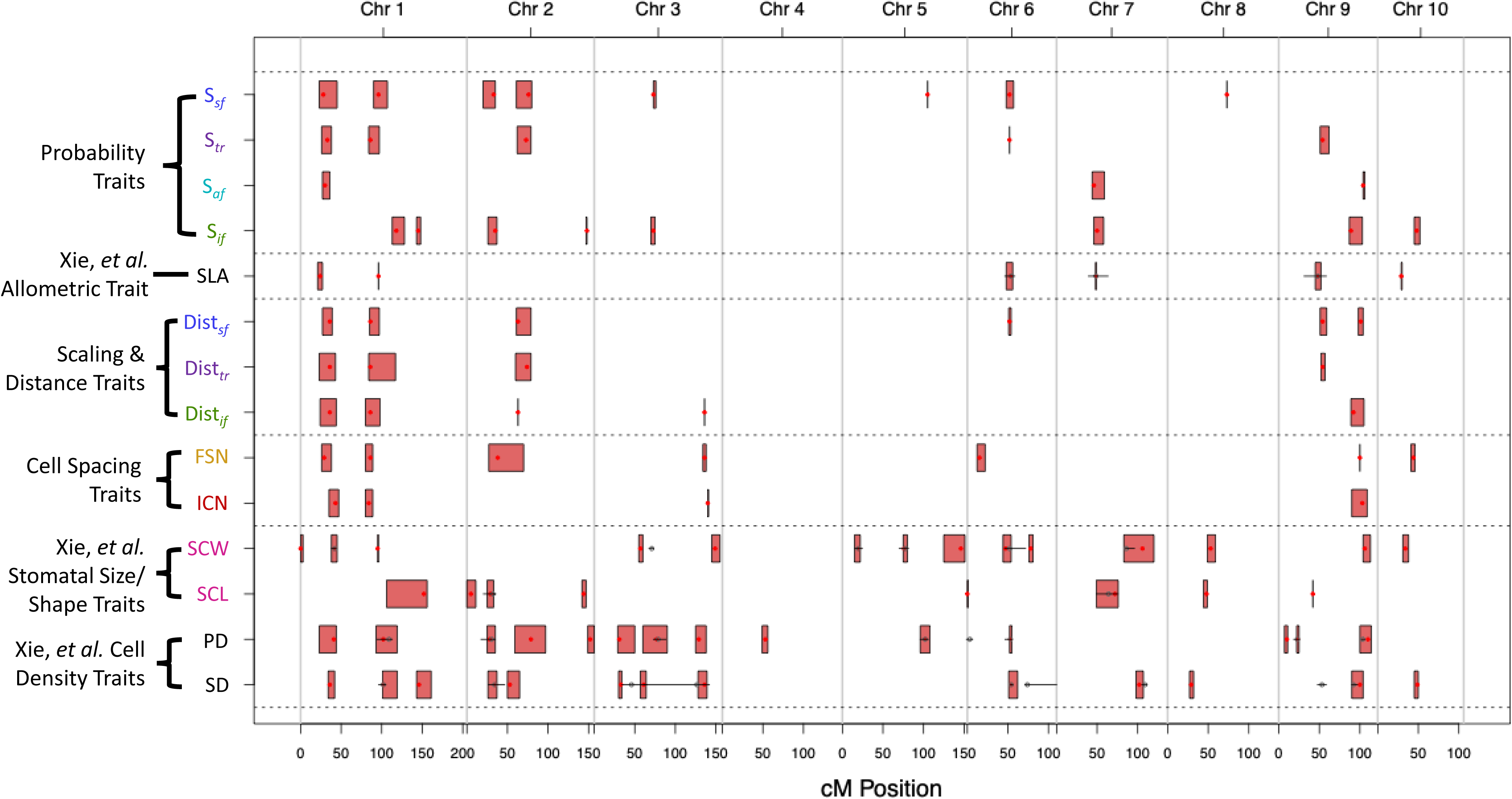
QTL intervals from multiple-locus scans of assorted Mask R-CNN and SPP traits for the 2016 field season (red boxes) and maximum LODs of each interval (red point). Previously identified QTL intervals identified by Xie are shown as hollow black points with whiskers corresponding to the QTL interval.

Of the thirteen SD QTL, twelve colocalized with QTL for the SPP traits **(Supp. Table 1)**. The two strongest SD intervals were QTL9 (LOD = 27.4, *PVE*=*16.3 %*) on chromosome 9 at 100 centimorgans (cM) and QTL1B (LOD = 19.7, *PVE*=*9.3 %*) on chromosomes 1 at 102 cM (**Fig. 6; Supp. Fig. 8A; Supp. Table 1**). These SD QTL were found to colocalize with various combinations of SPP traits. For example QTL1B colocalized with *S*_*if*_ (LOD=11.1, *PVE*=*7%*) and *S*_*sf*_ loci (LOD=8.8, *PVE*=*5.5%*). Meanwhile, *S*_*if*_ (LOD=17.9, *PVE*=*11%*) and ICN loci (LOD = 10.4, *PVE*=*6.6%*) colocalized with QTL9 (**Fig. 6; Supp. Fig. 8A, K, M, P; Supp. Table 1**).

QTL1A on chromosome 1 at 36cM was the third strongest SD interval (LOD = 15, *PVE*=*9.4 %*) and colocalized with *S*_*tr*_ (LOD = 26.6, *PVE*=*16%*) and *Dist*_*sf*_ loci (LOD = 25.5, *PVE*=*15.3%*) (**Fig. 6; Supp. Fig. 8A, J, O; Supp. Table 1**). In multiple cases, the LOD scores and PVEs for SPP trait QTL exceeded those exceeding their overlapping SD loci **(Supp. Table 1, *green LOD/PVE values*)**. Novel QTL associations for SPP traits were also found at loci previously only identified as SD QTL by Xie et al. (2021). For example, a minor SD QTL on chromosome 10 at 49cM colocalized with the *S*_*if*_ (LOD=6.3, *4%*) and FSN (LOD=6.2, *PVE*=*3.9%*) **(Fig. 6; Supp. Fig. 8A, L, M; Supp. Table 1)**. Despite these clear patterns of colocalization between SD and SPP traits, several SPP traits were only moderately heritable (**Supp. Table 2**). Although the SPP trait with the lowest narrow-sense heritability score, ICN (ℎ^2^ =0.196), still exceeded the heritability for SLA (ℎ^2^ =0.138).

## Discussion

This study successfully addressed its aims to exploit ML-generated HTP data to define and quantify a set of traits that describe spatial variation in stomatal patterning, and then discover the network of trait relationships and QTLs that underlie variation in SD across a maize RIL population. In doing so, it introduces a new spatial analysis method that can be applied universally to quantitatively describe stomatal patterning within and across grass species. It also provides a case study demonstrating how a comprehensive statistical network of component traits extracted from image data can be used to provide novel insights into the genetic basis for variation in a composite trait such as SD. This “deep phenotyping” approach is notably distinct from common approaches to investigation of trait interactions and quantitative genetics, where only a portion of the component traits needed to understand variation in a functionally important composite trait are measured and analyzed.

### The Core SPP traits Underlying Stomatal Density in the Maize B73 x MS71 RILs

The literature on grass leaf development suggests the cell file system should be a dominant source of SPP variation (McKown & Bergmann 2020; Nunes *et al*., 2020). This was substantiated by the spatial probability maps showing likely positions of neighboring stomata, with two prominent longitudinal peaks (i.e., the in-file neighbors) and the two minor lateral peaks (i.e. stomata in neighboring files; **Fig. 2**). *S*_*if*_ acted as a proxy measure for the stability of longitudinal runs of stomatal files and exhibited a correspondingly strong, positive relationship with SD throughout this study (**Fig. 2**, **Fig. 4**, **Fig. 5**, **Fig. 6**). This was most recognizable in the longitudinally flattened SPP histograms (**Supp. Fig. 3B**) where a clear gradient of high to low SD values (gold to purple) appeared at the origin (**Supp. Fig. 3C-H**). The SPP morphology was generalizable across the RILs, indicating that it is a conserved, self-repeating, patterning scheme for the local stomatal ‘neighborhood’ in maize. Although the variances around these peaks was anticipated to be important for describing density, preliminary analyses suggested that these variances did not carry significant information beyond that in their respective spacing and distance measures (**Fig. 3**, **Supp. Fig. 5**). Increasing distances between neighboring stomata by itself should be sufficient to increase disorder in the distribution of probabilities described by the SPP. This is illustrated by the lowest SD genotype, Z019E0140, having the greatest centroid distances as well as the most diffuse probability map (**Fig. 2H**). Given there were clear signs of trait co-correlations it was necessary to view this phenotype as a structured system.

### Modeling Informs which SPP traits Predict SD and their Structural Interactions in the B73 x MS71 RILs

Initial evidence indicated the SPP traits were often co-correlated (**Fig. 4**). PCA and comparison of linear models accounting for different combinations of traits were able to identify a core set of traits that explained SD (**Supp. Fig. 5,6**). Piecewise structural equation modeling (SEM) of these core traits allowed us to explore their causal interactions (**Fig. 5**). The SEM results indicated a hierarchy beginning with cell size and spacing traits with SCL and ICN defining longitudinal effects and SCW and FSN for similar lateral effects (**Fig. 5**). These initial traits feed variation into the subsequent triad of probability phenotypes (*S*_*if*_, *S*_*tr*_, *S*_*sf*_) that define the distribution of the SPP itself, and ultimately SD. First in this hierarchy of probability traits is *S*_*tr*_ and then *S*_*sf*_, with both feeding their variation into the *S*_*if*_that subsequently is the largest influence on SD in this RIL population (**Fig. 5**). The strongest path observed in the SEM is FSN→ *S*_*tr*_ → *S*_*if*_ →SD with the first two traits potentially being related to vascular patterning and even broader leaf allometric relations (Al-Salman *et al*., 2023; Facette *et al*., 2015; McKown & Bergmann 2020; Nunes *et al*., 2020). Evidence from the primary literature generally agrees that grass stomatal files form immediately adjacent to veins potentially due to their integrated roles in regulating plant water status and CO_2_ diffusion distance (Holbrook & Zwienicki, 2005; Niklas, 1994).

Broadleaf systems are typically more mesophyll dominated, exhibiting stronger association between SLA and mesophyll anatomy, particularly palisade parenchyma formation (John *et al.,* 2017). Grasses by contrast exhibit a more homogenized leaf internal anatomy which is further exacerbated in C4s (e.g., maize and sorghum) having a higher vein density (Al-Salman *et al*., 2023; Kumar & Kellogg, 2018). C3 grasses (e.g., Brachypodium) could be anticipated to exhibit multiple epidermal files between veins dampening this SPP trench/stomatal file system/SLA association, a possibility well worth exploring in future studies of this phenotype (McKown & Bergmann 2020; Nunes *et al*., 2020). The higher minor vein densities common in this C4 context of maize likely only provides enough space for a single stomatal file to be equidistant between veins flanking them on either side (Sedelnikova *et al*., 2018). This flanking vein morphology effectively creates the boundary between stomatal files we observe as the SPP trench and would also help explain the associations observed between *S*_*tr*_/*S*_*sf*_ and SLA (**Fig. 4 & 6, Supp. Fig. 5**; Kumar & Kellogg, 2018). Despite elements of this path analysis agreeing with the current literature, these hierarchies should not be conflated with developmental sequence. For example, SCL occurs early within the SEM hierarchy, yet it is likely the product largely of expansion growth and should be one of the last traits to arise developmentally (i.e., after all key cell divisions like ICN have transpired) based on the sequence of events described in the literature and the mechanical links between cell wall remodeling, turgor and expansion growth (McKown & Bergmann, 2020; Nelissen *et al*., 2012; Saibo *et al*., 2006). While imperfect, aspects of this SEM do appear to mirror what is known about development, with lateral traits like FSN, *S*_*tr*_, and *S*_*sf*_ reinforcing the variation observed in *S*_*if*_ which then acts as a nexus node that drives SD (**Fig. 5**). Despite the importance of in-file variation in explaining SD in the SEM, its precursors SCL and particularly ICN, have less prominent roles than anticipated (**Fig. 5**). Intervening pavement cells typically arise from the initial asymmetric divisions that also canalize guard mother cells so this lack of variation could be a more fundamental limit for stomatagenesis, at least in the context of the present population (Asraf *et al*., 2023; Facette *et al*., 2015; Lunn *et al*., 2024, Zhou *et al*., 2025). Leveraging populations with greater genetic diversity in future studies will also allow for more critical assessments of whether limited genetic variation for key patterning measures such as ICN is a more general phenomena or simply due to the alleles available in Ms71 and B73. In time, it will also worth exploring this phenotype in other model systems to see how well these core SPP traits are conserved in other grass systems or potentially even dicots.

### Segmenting the Genetic Underpinnings of SD into its Potential SPP Trait Influences

Stomatal patterning has been found to be polygenic across numerous plant systems (Delgado *et al*., 2019; Ferguson *et al*., 2021; Gailing *et al*., 2008; Xie *et al*., 2021). Despite the capacity to identify multiple stomatal density, size, and shape loci (traits that are often quite heritable) the component traits underlying our SPP phenotype, particularly ICN, appeared to be only modestly heritable by contrast (**Supp. Table 2**). There are three competing interpretations for this discrepancy, although the latter two seem the most likely: (1) the phenotypic variances for these phenotypes are essentially random, (2) the genetic variance available within this RIL population may be limited, and lastly (3) some of these component traits may not necessarily behave in an additive manner (Templeton, 2006). As noted above, even though the average stomatal pattern is highly predictable within a genotype, it is still probabilistic, either due to factors of broader leaf anatomy which may impact adjacent stomatal files, or even random canalization failures which produce guard mother cells (Ashraf *et al*., 2023). A second interpretation is that the key developmental processes under study could be genetically constrained, only bearing weak alleles, if any, in natural populations. This would reflect a strong selective pressure to maintain wild-type function that would consequently lead to there being little heritable variation to recognize (Hansen *et al*., 2011; Templeton, 2006). The discrepancy between the narrow and broad sense heritability presents another possibility however as the former largely captures additive (i.e. linear) effects alone, although the broad sense suggests that the underlying genetic basis of these traits could potentially be more complex.

Like the previous investigation using this population (Xie et al. 2021), SD was found to be polygenic. However, the current investigation was able to glean novel spatio-developmental insights into what were previously described as generic SD loci (Xie et al., 2021). SD loci typically associated with one or more component trait QTL in combinations that were often indicative of particular roles in stomatal/leaf development (**Fig. 6, Supp. Table 1**). Thus, the SPP approach allowed a number of QTL to be re-categorized as being QTL for specifically longitudinal versus lateral components of SD. Most strikingly, the first and third most prominent SD loci, QTL9 and QTL1A, were re-defined as loci underlying longitudinal and lateral components of stomatal patterning, respectively (**Supp. Table 1**). This capacity to interrogate how SD variation might have arisen from distinct elements of stomatal patterning, which likely occur at different times in leaf development can guide more focused follow-up studies.

Given the direct association between *S*_*if*_ and SD in the SEM, it was unsurprising that the strongest effect SD locus in this study, QTL9, appeared to act through this *S*_*if*_ → SD pathway (**Fig. 5**, **6**). This ICN or *S*_*if*_ route was the most immediate route to alter SD in the SEM (**Fig. 5**). As noted above there is a concern of severe developmental defects in modifying ICN or *S*_*if*_making “milder” effect alleles like QTL9 a clear target for future research. This might suggest that although genetic diversity for longitudinal spacing factors may be limited, at least in this population, (discussed above) this variation may still provide a more immediate route to dramatically altering SD. Despite the absence of pavement cell information with the SPP, colocalization of SPP traits and pavement cell density (PD) from Xie et al. (2021) suggests that the SPP approach does capture a holistic view of epidermal patterning from analysis of a single cell type.

### The Novelty and Implications of the Grass Stomatal Patterning Phenotype (SPP)

Stomata are one of the best studied plant cell types because of their accessibility on the epidermis, direct links to developmental biology and cell canalization, as well as their importance to leaf physiology, plant productivity and resilience (Baresch *et al*., 2019; Lawson and Leakey 2024; Nunes *et al*., 2020). Stomatal density has been a trait of particular focus because it directly impacts the capacity for the leaf to exchange gases with the atmosphere, and it is impacted by the frequency with which epidermal cells acquire stomatal identity (Raissig *et al*., 2016; Zhou *et al*., 2025). Spatially explicit developmental processes have been known to drive variation in stomatal density, but most studies have still been limited to intuiting these processes from the census measures SD or SI (Casson & Gray, 2008; Doheny-Adams *et al*., 2012; McKown & Bergmann 2020; Nunes *et al*., 2020). This reflects the much greater effort required to collect data on stomatal patterning than stomatal density, and the challenge of quantifying spatially explicit measures of stomatal patterning that are generalizable across the diversity of epidermal anatomy found in plants (Conklin *et al*., 2018; Vőfély *et al*., 2018).

Although SD is the result of spatial processes it cannot by itself describe these multidimensional sources of variation. For example, increasing the number of pavement cells separating stomata within a file of cells, or increasing lateral spacing with more files of pavement cells separating files of cells containing stomata, would both reduce stomatal density, but by different developmental processes regulated by different sets of genes (**Fig. 7**). In addition, an orthogonal source of variation in stomatal patterning can come from differences in cell size, which works to alter spacing and stomatal density independent of any changes in cell identity (**Fig. 7A-B**). Normalizing the output of nearest neighbor analysis for cell size allows the SPP approach to estimate key spacing information (**Fig. 7C-F**). The network of SPP traits provides a novel and efficient means to rapidly extract a holistic set of spatial features that govern SD in grasses (**Fig. 1**). Many of these SPP traits were co-correlated (**Fig. 4**, **Supp. Fig. 5**) allowing for a simplified set of core SPP traits which generally could be classified into: (1) cell size, (2) cell packing, and (3) patterning probability categories, which collectively explain SD.

**Figure 7.**
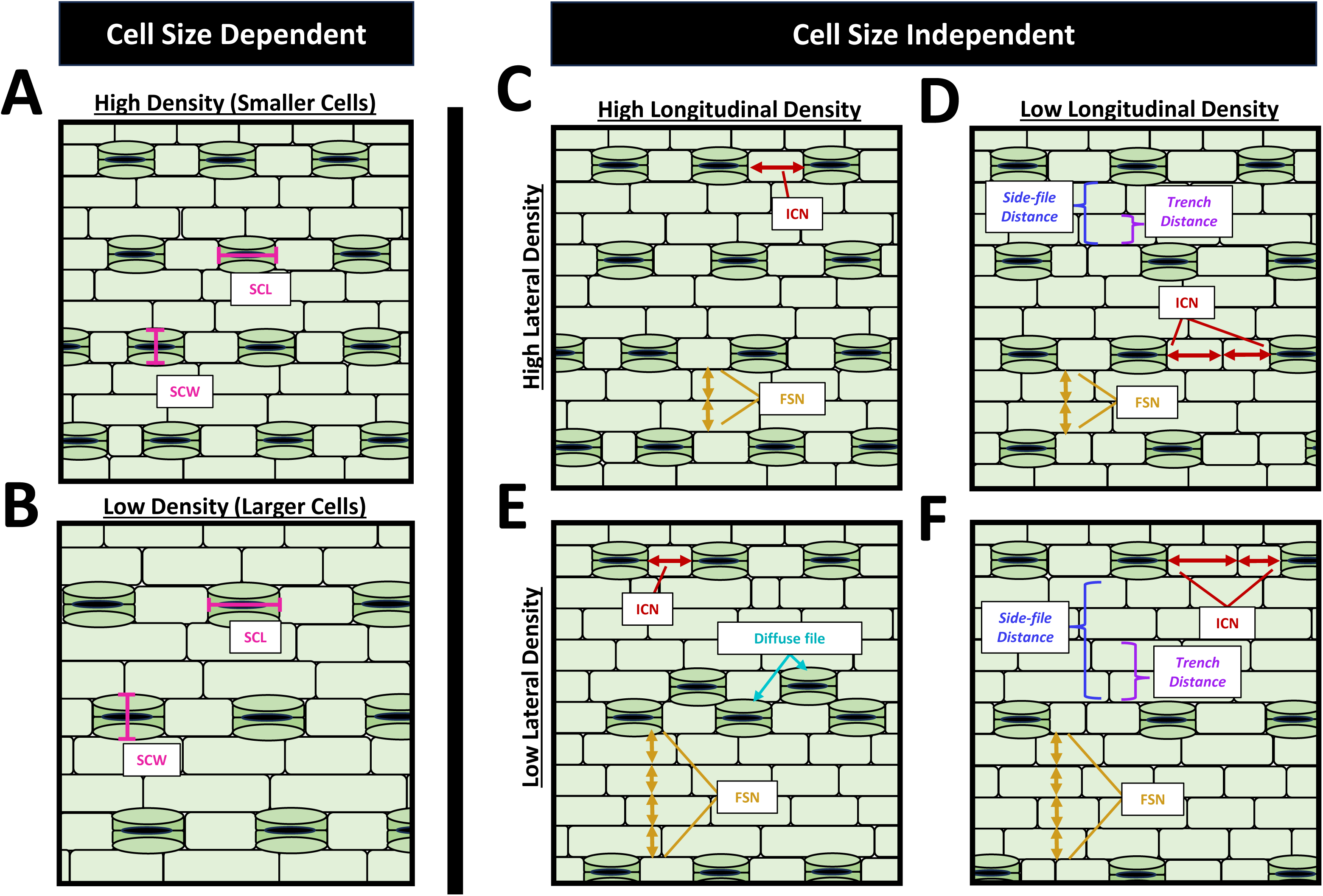
Overview of the core traits that can drive variation in the stomatal patterning phenotype which consist of two clear categories cell size dependent traits **(A-B)** and cell size independent traits **(C-F)**. Cell size which can be impacted by SCL and SCW is likely a key feature in stomatal patterning and SD overall with smaller cells **(A)** allowing for a denser arrangement of stomata whereas increasing cell size **(B)** would inversely lower density by creating larger pores. Stomatal density can also be amended through cell packing rather than cell size variation, for example high stomatal densities **(C)** can be lowered either by increasing longitudinal spacing via ICN **(D)** or through increasing lateral spacing via FSN **(E)**, and by increasing spacing in both directions **(F)** decreases in SD should in principle be maximized.

Stomatal patterning can be found in particularly diverse forms across the dicots, which includes many of the species that have been the dominant model systems for molecular genetic research i.e. Arabidopsis, poplar, tobacco, and tomato. By contrast, the genes controlling stomatal development in grasses have been significantly less studied until recently, despite the more constrained arrangement of epidermal cell patterning that they possess (Doll *et al*., 2023; Spiegelhalder & Raissig *et al*., 2021; Torii, 2021). ML-enabled computer vision analysis of epidermal micrographs for high throughput phenotyping of stomatal patterning has recently emerged for many species (Tan et al. 2024). This study exploited the tractability of grass stomatal patterning to demonstrate how spatial analysis of the output from such tools can be mined to extract greater information value.

The SPP approach leverages the fact that the spatial arrangement of stomata in a mature leaf relates to series of distinct histogenic events that occurred while it was developing (McKown & Bergmann 2020; Nunes *et al*., 2020). For example, there is mounting evidence for relationships between internal vasculature (established early in leaf development) and subsequent epidermal stomatal file patterning (Facette *et al*., 2015; McKown & Bergmann 2020; Nunes *et al*., 2020). Therefore, when the SPP approach allows QTL that were previously generically identified for SD to be redefined as specific to traits that relate to the lateral versus longitudinal components of SD, it suggests which phases of leaf development the underlying genes are likely to have been active in. Mammalian pad ridge morphologies (fingerprints) may offer a developmental genetic parallel for epidermal characters associated with internal anatomy. Like stomatal patterning, pad ridges result from earlier internal developmental influences (i.e., volar pad size) whose existence is largely limited to early fetal limb development (Li *et al*., 2022). Co-correlations between traits are more intuitive when viewed from this perspective since leaf histogenesis, venation and stomatal patterning (much like volar pad and subsequent digit and fingerprint development) may be indicative of a developmentally integrated organ system.

The SPP trait approach has two features that help make it translatable to the broader study of stomatal patterning, First, the SPP method provides a conceptual framework that encapsulates stomatal patterning as a self-repeating pattern with a few key spatial elements related to longitudinal or lateral aspects of grass leaf morphology. Second, it is robust even though stomatal patterning is heterogeneous at several scales e.g. stochasticity in the identity of cells immediately neighboring one another, as well as systematic variation in patterning among locations at different lateral transect distances from the midvein. As a result, the typical grass morphology should make the SPP approach applicable to all other grass species (Devos & Gale 2000; Doust & Diao, 2017; Garvin *et al*., 2008; Mullet *et al*., 2014).

## Materials & Methods

### Machine Learning Annotated Maize Leaf Data

3785 micrographs, each showing ∼800 *μm*^2^ of the abaxial surface of the youngest fully expanded leaf (**Fig. 1A-B**), from 180 lines of a B73 x Ms71 RIL population of maize grown during the 2016 field season were previously collected and the locations and sizes of stomata determined using a Mask R-CNN workflow (Xie *et al*. 2021). For the current study, the coordinates of the centroid of each stomatal mask and their size (length and width) was analyzed. The x-dimension corresponded to the longitudinal axis of the leaf (i.e. along the length of the leaf blade) and the y-axis corresponded to the lateral axis of the leaf (i.e. from midvein to leaf margin).

### Overview of Stomatal Patterning Phenotype (SPP) analysis pipeline

Quantification of the SPP involved: (1) assessing nearest neighbor distances and positions between stomata (**Fig. 1C-E**); (2) normalizing the nearest neighbor distance and positional information with respect to the origin so as to center them around a common “origin stomata” (**Fig. 1F**); (3) generalizing this spatial information from these normalized scatterplots into 2-dimensional kernel density distributions (i.e., a 2-dimensional probability distribution; (**Fig. 1G**); and (4) extraction of traits quantifying the variation observed in this spatial phenotype (**Fig. 3**. The initial workflow described in 1-4 and the necessary functions to perform them on this dataset form the Python SPP library available on Github (https://github.com/jgerardhodge/pySPP). Subsequent analyses were performed in R to: (5) Identify the smallest subset of traits that collectively describe the variance in SD i.e. dimension reduction via correlation, principal component analysis (PCA), and linear modeling predictions of SD; (6) build a structural equation model (SEM) to determine the network of trait interactions underlying variation in SD, and (7) QTL mapping to assess how genetic effects spatially segregate.

### Ranked Nearest Neighbor Approximate of the Average Stomatal Neighborhood

The *stomata_rankedNN* function calculated the nearest neighbor distance, “*i*”, between the centroid of an origin stomata and a neighboring stomata for all “1…*n*” stomata within the image using the following Manhattan distance formula (Reynolds, 1980):

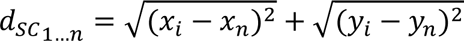

Manhattan distances are measured along stair-step patterns in cardinal directions, which is appropriate to quantification of the linear, tiled arrangement of epidermal cells on the leaf epidermis of grasses (**Fig. 1C-E**). To avoid edge effects, only stomata that were at least 50 *μm* away from the edge of the image were allowed to act as origin stomata (**Fig. 1C-E, dotted grey box**). The distance and direction to the five nearest neighbors were then expressed on a relative basis for each origin stomata (**Fig. 1F**).

### Kernel Density Distributions (KDDs) of the Stomatal Patterning Phenotype (SPP)

The spatial distribution of normalized nearest neighbors varied among genotypes, but with a shared geometric structure (**Fig. 3A-D**). This means the method is generalizable and phenotypic variation could be quantified as a 2-dimensional kernel density distribution using the *stomataKDDs* function at a 1 *μm*^2^ kernel resolution over an area ±150 *μm* bounds from the origin (**Fig. 3E-H**). Across the entire RIL population, only 8 out of the115,537 stomata identified as a 5^th^ ranked nearest neighbor exceeded these 150 *μm* bounds (**Supp. Fig. 9**). Depending on the feature of stomatal patterning being assessed, the kernel density distribution was produced for different combinations, or an average of all, of the 1^st^ to 5^th^ ranked nearest neighbor stomata.

### SPP Trait Annotation and Extraction

Four distinct hotspots were defined by the SPP that describe the local neighborhood around stomata. Nearest neighbors were either found within the same file of epidermal cells (i.e. horizontally on the image and longitudinally on the leaf, so termed “in-file hotspots”) or they could represent stomata in flanking files of cells (i.e. vertically on the image and laterally on the leaf, so termed “side-file hotspots”). These in-file and side-file hotspots could be readily identified in the probability distributions of all 1^st^-5^th^ nearest neighbor stomata for any given RIL (**Fig. 2E-H**). The in-file and side-file hotspots were separated by a pronounced “probability trench” where stomata tended not to occur.

A series of 15 SPP traits (**Table 1**) were identified that characterize the position, magnitude, variation and relative proportions of these zones of high/low probability, along with measures of stomatal complex size (**Fig. 3**).

**Table 1.** Index of traits used in this investigation that were either previously generated by Xie, et al. using Mask R-CNN or are extracted SPP phenotypes with corresponding abbreviations, descriptions, and calculations used to generate them. § denotes the trait identifying genotypes that do or do not possess diagonally adjacent stomata. * denotes traits identified as core descriptors of stomatal patterning.

| Trait Abbreviation | Category | Data source used to estimate trait | Trait Description | Estimation Method |
| --- | --- | --- | --- | --- |
| *SD* * | Cell/complex density | Mask R-CNN analysis of epidermal cells in field-of-view | Stomatal complex density |  |
| *PD* |  |  | Pavement cell density |  |
| *SCL* * | Stomatal complex morphology |  | Stomatal complex length |  |
| *SCW* * |  |  | Stomatal complex width |  |
| *SLA* | Allometry | Destructive harvest | Specific leaf area for the given leaf sample |  |
| *S<sub>if</sub>* * | Probability intensity | Probability map of the 5 nearest neighboring stomata | In-file probability = magnitude of the peak in probability for stomata in the immediate neighborhood being in the same cell file as the origin stoma (Fig. 2a) |  |
| *S<sub>tr</sub>* * | | | Trench probability = magnitude of the minimum in probability for stomata in the immediate neighborhood being at positions immediately lateral to the origin stoma (Fig. 2a) | $\frac{\text{Upper } S_{tr} + \text{Lower } S_{tr}}{2}$ |
| *S<sub>sf</sub>* * | | | Side-file probability = magnitude of the peak in probability for stomata in the immediate neighborhood being at positions lateral to the origin stoma (Fig. 2a) | $\frac{\text{Upper } S_{sf} + \text{Lower } S_{sf}}{2}$ |
| *S<sub>af</sub>*§ | | Probability map of the 1st nearest neighboring stoma | Diffuse file probability = magnitude of the probability that the 1<sup>st</sup> nearest neighbor is diagonally adjacent to the origin stoma (Fig. 2b,c) | $\frac{\text{Upper } S_{tf} + \text{Lower } S_{tf}}{2}$ |
| *Dist<sub>tr</sub>* | Distance among probability hotspots | Probability map of the 5 nearest neighboring stomata | Distance between the lateral trench and the origin stoma (Fig. 2a) |  |
| *Dist<sub>if</sub>* |  | Probability masks of the 1<sup>st</sup> and 2<sup>nd</sup> nearest neighboring stomata | Distance between the In-File peak mask centroid and the origin (Fig. 2f) |  |
| *Dist<sub>sf</sub>* |  | Probability masks of the 4<sup>th</sup> and 5<sup>th</sup> nearest neighboring stomata | Distance between the Side-File peak mask centroid and the origin (Fig. 2g). |  |
| *S<sub>ifLongiVar</sub>* | Variances/shape among probability hotspots | Probability masks of the 1<sup>st</sup> and 2<sup>nd</sup> nearest neighboring stomata | Bounding box length estimate of longitudinal variance of the In-file peak mask (Fig. 2f). |  |
| *S<sub>ifLatVar</sub>* |  | Probability masks of the 1<sup>st</sup> and 2<sup>nd</sup> nearest neighboring stomata | Bounding box width estimate of lateral variance of the In-file peak mask (Fig. 2f). |  |
| *S<sub>ifVarArea</sub>* |  | Probability masks of the 1<sup>st</sup> and 2<sup>nd</sup> nearest neighboring stomata | Bounding box area estimate of spatial variance of the In-file peak mask (Fig. 2f). |  |
| *S<sub>sfLongiVar</sub>* |  | Probability masks of the 4<sup>th</sup> and 5<sup>th</sup> nearest neighboring stomata | Bounding box length estimate of longitudinal variance of the Side-file peak mask (Fig. 2g). |  |
| *S<sub>sfLatVar</sub>* |  | Probability masks of the 4<sup>th</sup> and 5<sup>th</sup> nearest neighboring stomata | Bounding box width estimate of lateral variance of the Side-file peak mask (Fig. 2g). |  |
| *S<sub>sfVarArea</sub>* |  | Probability masks of the 4<sup>th</sup> and 5<sup>th</sup> nearest neighboring stomata | Bounding box area estimate of spatial variance of the Side-file peak mask (Fig. 2g). |  |
| *ICN* | Cell or file spacing | Probability mask of the 1<sup>st</sup> and 2<sup>nd</sup> nearest neighboring stomata | Estimated intervening pavement cell number between In-file stomatal complex neighbors (Fig. 2f). | $(\frac{d_{IF}}{SCL} - 1)$ |
| *FSN* | | Probability mask of the 4<sup>th</sup> and 5<sup>th</sup> nearest neighboring stomata | Estimated pavement file number acting as spacers between stomatal files (Fig. 2g). | $(\frac{d_{SF}}{SCW} - 1)$ |

A set of 4 traits describing spatially distinct “probability intensities” were annotated by: (1) averaging the probabilities within horizontal rows of pixels of each SPP, effectively flattening the heatmap into a vertical histogram; then (2) extracting the probabilities corresponding to in-file (*S*_*if*_) and side-file (*S*_*sf*_) peaks and trenches (*S*_*tr*_) in the profile (**Fig. 3A**) as defined in Table 1. Although not used for trait annotations in this study, an orthogonal operation was also performed by averaging the probabilities within vertical columns of pixels of each SPP to produce horizontal histograms. Notably, diffuse in-file peaks were found in some genotypes in which two neighboring files shared a stomatal identity. This was easiest to assess as a adjacent-file probability (*S*_*af*_) one cell width way from a SPP vertical histogram of only the 1^st^ nearest neighbors, while all other probability traits in this study are assessed from the average of the 1^st^-5^th^ nearest neighbor SPPs (**Fig. 3B, C**).

A set of 3 distance traits were annotated by measuring either the centroid distance between the two primary hotspot classes (i.e. the in-files and side files), or the mid-position of the lateral trenches to the origin stoma (**Fig. 3D-G**), as defined in Table 1. In-file SPPs were generated using only the 1^st^ and 2^nd^ ranked nearest neighbors to assess in-file distances, while 4^th^ and 5^th^ ranked nearest neighbors were used to generate comparable side-file SPPs for assessment of distances among lateral features (**Fig. 3D, E**). In both cases, the SPPs were thresholded at their 55^th^ percentile to generate binary masks prior to trait estimation (**Fig. 3F, G**). Given that the positional information of the trenches was most apparent in the vertical probability histograms described above the trench distances were typically measured alongside their *S*_*tr*_ intensity measures (**Fig. 3A**).

A set of 6 “peak spatial variances” were also annotated alongside the above-mentioned in-file and side-file peak distances by applying bounding boxes to these peak masks which allowed for longitudinal variances to be defined for the in-file (*S*_if _*LongiVar*__) and side-file (*S_if *LongiVar*_*) peaks (**Fig. 3D-G**). Corresponding lateral variances for the in-file (*S_if *Long*.*Var*_*) and side-file (*S_sf *Long*.*Var*_*) peaks as well as the variance areas of the in-file (*S_sf *LongiVar*_*) and side-file (*S_if *Long*.*Var*_*) peaks were also annotated (**Fig. 3D-G**). A set of 2 “cell/file spacing” traits were generated to estimate the amount of cell packing within the longitudinal and lateral orientations (**Fig. 3D-G**), as mathematically defined in Table 1. The intervening cell number (ICN) estimated the number of pavement cells between successive stomata in a single file (**Fig. 3F**). Meanwhile, the pavement file spacing number (FSN) estimated the number of nonstomatal files of cells between an origin stoma and its nearest laterally located neighboring stomatal file (**Fig. 3G**). These calculations assumed stomatal complex width and length to be rough proxies for pavement cell size in their respective orientations.

### Trait Dimensionality Reduction and Associations

Principal Component Analysis (PCA) and Pearson trait correlations were leveraged on the full trait dataset (Table 1) to select a series of candidate trait sets that captured a large portion of the variance in this ordination space with the minimal number of traits (**Supp. Fig. 5**). These trait-trait relationships were complemented by pairwise trait relationships of Pearson correlation coefficients, generated with the base R function *cor*, which were visualized using the R function *corrplot* within the *corrplot* library (Wei & Simko, 2021). A series of SD linear model predictions were then used to further assess which traits must be included in a set of core SPP traits (**Supp. Fig. 6**). The underlying functional hierarchy among the core set of seven traits was assessed using piecewise structural equation models (pSEMs) (Shipley, 2000, Lefcheck, 2015). The pSEMs were iteratively optimized based on the Akaike Information Criterion (AIC) statistics of each model iteration as well as tests of directed separation to resolve missing trait-trait relationships within each model generation. These operations were performed using base R’s *lm* and *predict* functions in addition to the *psem* function found in the *piecewiseSEM* library (Lefcheck, 2015).

### Genetic Mapping of SPP Traits

All traits were merged with an existing genotype-by-sequencing (GBS) map (Xie et al. 2021) following the *R/qtl* genetic map format and imported into R using the qtl function, *read.cross* and the *convert2riself* function. The data was converted to a *qtl2* format using the R library *qtl2*’s function *convert2cross2* (Broman, *et al*. 2019). To provide an even distribution of markers within each linkage group a minimal number of pseudomarkers were generated for this RIL map design using the *insert_pseudomarkers* function. Genotypic probabilities for this map were calculated using the *calc_genoprob* function with *error_prob=0.002*. A more stringent LOD threshold of *α*=0.0005 (allowing focus on the loci of largest effect) was dynamically generated for each trait using the *scan1perm* function to perform 1000 permutations of each trait measure. Multi-locus QTL scans were performed for each trait via a Haley-Knott regression using the *scan1* function. Intervals exceeding LOD scores of the dynamic *α* threshold were filtered with the max LOD marker being taken as the location of associated QTL for each trait. Additionally, for multiple QTL to be detected with this function the peak drop difference (i.e., the amount LODs must decrease between peaks in a continuous region greater than the LOD threshold) was set to the dynamic *α* threshold to ensure stringency. Finally, the edges of each QTL interval around these max LODs were defined as a drop in LOD score greater than 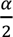 from the max LOD. In some cases, weak QTL could produce a max LOD greater than *α* while still having a 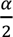 that was lower than the median LOD score present for the map, effectively creating erroneously large QTL intervals for weak effect regions. In these instances, the QTL intervals were replaced with those forming the margins for a continuous region surpassing the dynamic *α* threshold. Heritabilities were estimated using the *sommer* library in R, first by generating additive, dominance, and epistatic relationship matrices then leveraging this data with the mixed model function *mmer* to generate variance components for narrow and broad-sense calculations (Covarrubias-Pazaran, 2016).

## Acknowledgements

We would like to Kevin Xie for his assistance in leveraging his Mask R-CNN annotations while developing this workflow. In addition, we would also like to thank Elizabeth (Toby) Kellogg, Daniel Trejeda-Lunn, and James Fischer for their helpful discussions on the context of this spatial phenotype and its implications over the course of this investigation.

## Funding

This work was funded by the DOE Center for Advanced Bioenergy and Bioproducts Innovation (U.S. Department of Energy, Office of Science, Biological and Environmental Research Program under Award Number DE-SC0018420) and a generous gift from Tito’s Handmade Vodka. Any opinions, findings, and conclusions or recommendations expressed in this publication are those of the author(s) and do not necessarily reflect the views of the U.S. Department of Energy.

---

**Supplementary Figure 1.**
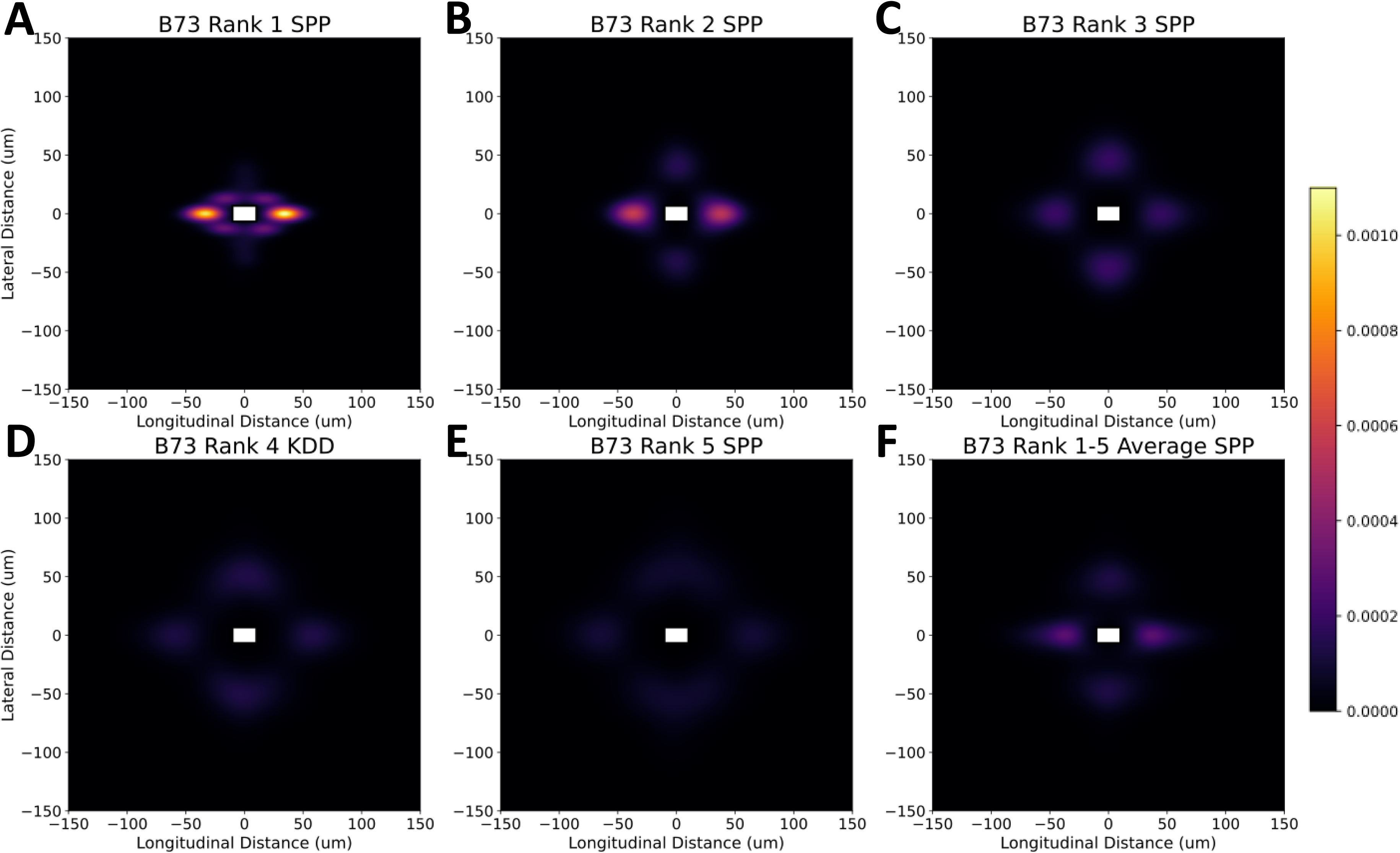
Variation underlying individual rank-ordered neighbors which are normalized to the same color map intensity ranges. When kernel densities are generated for specific nearest neighbor (NN) relationships as opposed to several averaged together a variation in how moduli are distributed around the origin stomata becomes more distinct. The immediate 1° **(A)** and 2° **(B)** rank-ordered neighbors’ character in-file neighbor relationships with the deltoid adjacent file moduli and probability trench between these vertically oriented neighbors and the origin stomata becoming more apparent in the 3° **(C)**, 4° **(D)**, and 5° **(E)** relationships although with higher orders the overall information does begin to diminish with each increasing rank. The average SPPs (F) are a product of these distinct layers being averaged at each kernel position. Supplementary Figure 5.

**Supplementary Figure 2.**
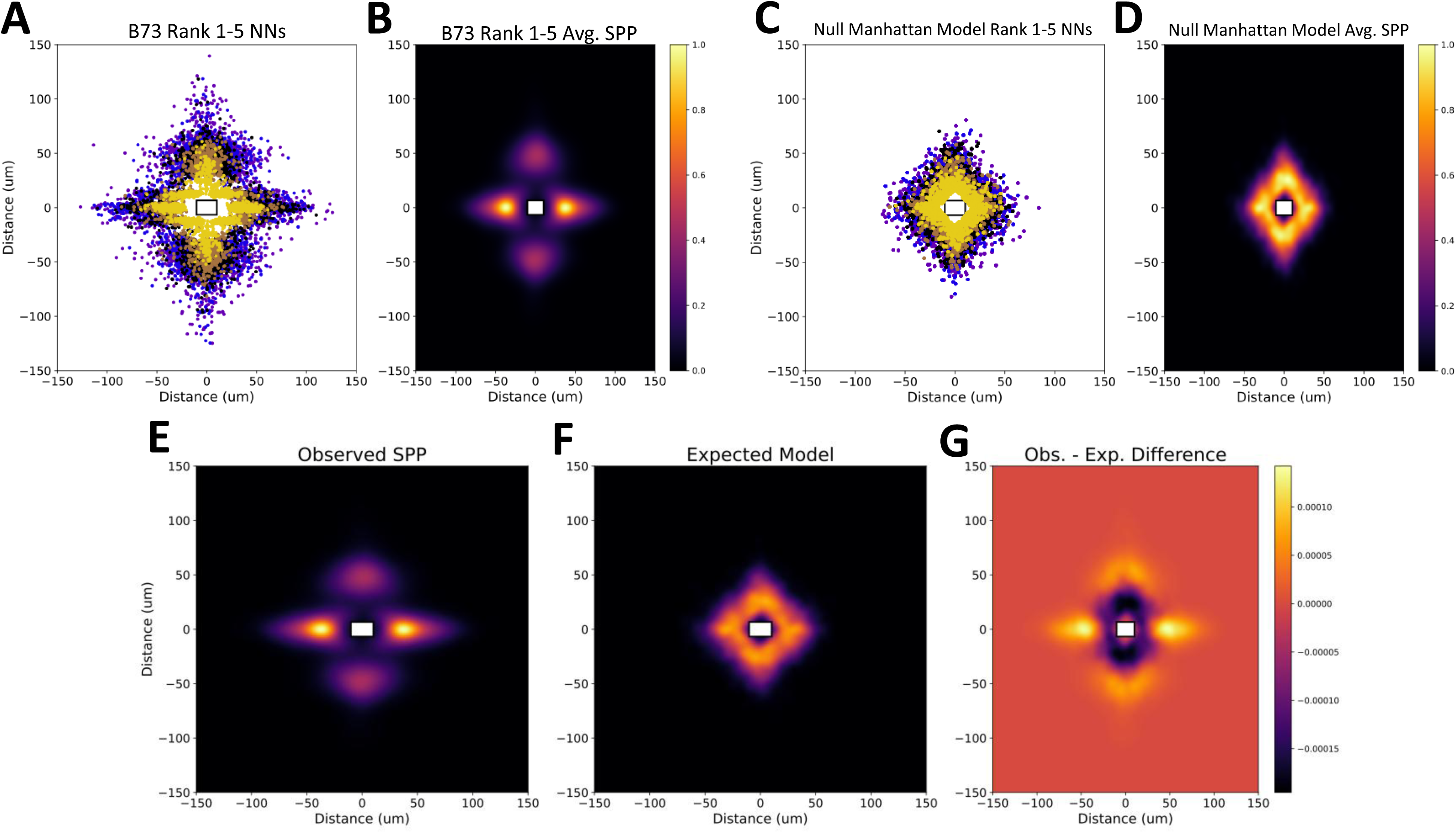
Validating the spatial patterns observed in the SPP phenotype are not artifacts of the Manhattan distance equation by comparing the NN observations from B73 to a null model. The observed data from B73 represented both as **(A)** an origin stomata scatterplot and **(B)** an SPP distribution. The null Manhattan distance model to compare against represented both as **(C)** an origin stomata scatterplot and **(D)** an SPP distribution. When the observed and null SPP’s **(B,D)** are normalized to the same scale **(E,F)** they can then be compared for their differences at each kernel position **(G)** in which a pronounced cold ring emerges where the Manhattan diamond is under represented in the observed data where as hot “tails” can be seen on the flanks of each corner, particularly the right and left, suggesting the null model is insufficient at capturing the tails around the observed in-file and side-file hotspots.

**Supplementary Figure 3.**
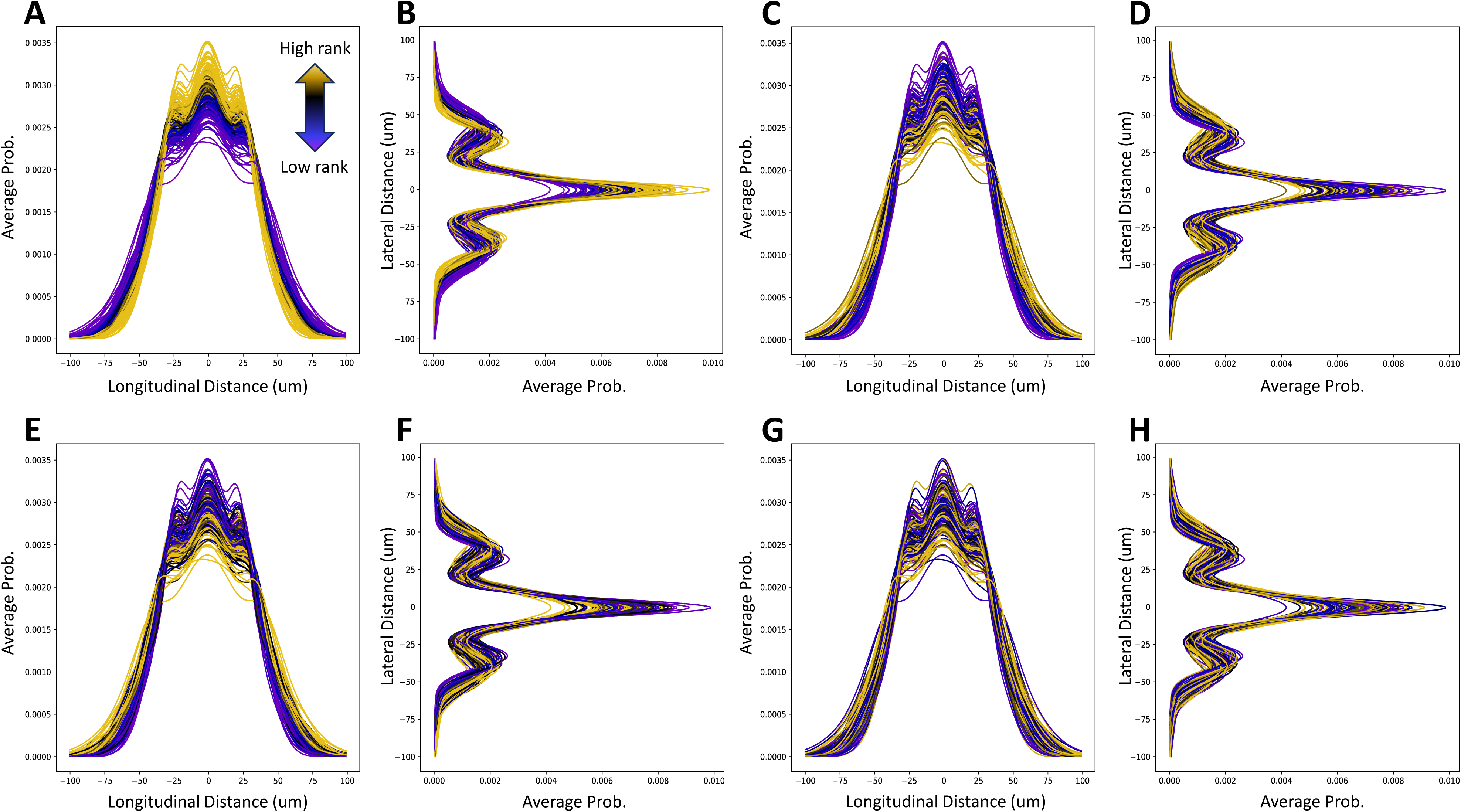
Relationships between flattened kernel density distributions and other stomatal traits. To allow for population wide comparison of kernel densities these 2D probabilities were flattened either horizontally **(A,C,E,G)** or vertically **(B,D,F,H)** to allow for all accessions within the RIL population to be plotted simultaneously. Furthermore rank-ordered values for **(A,B)** stomatal density (SD), **(C,D)** stomatal complex length (SCL), **(E,F)** stomatal complex width (SCW), and **(G,H)** stomatal complex area (SCA) could be color coded from minimum (purple) to maximum (gold) ranked values resulting in a gradient for stomatal density that is absent in other size related stomatal traits.

**Supplementary Figure 4.**
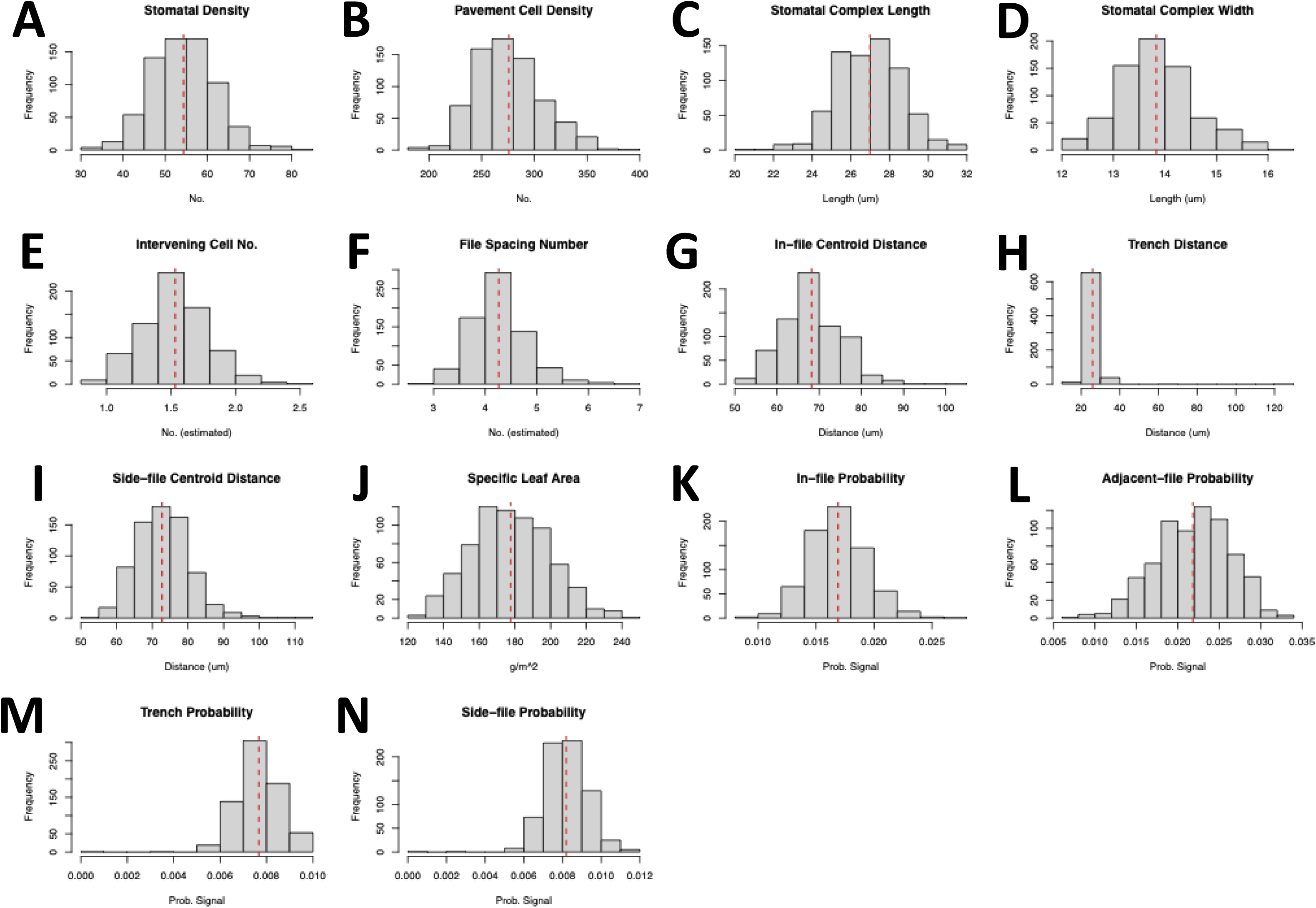
Histograms of the Xie *et al*. and SPP traits of interest leveraged in this study of the 2016 RIL population consisting of **(A)** SD, **(B)** PD, **(C)** SCL, **(D)** SCW, **(E)** ICN, **(F)** FSN, **(G)** *Dist_if_*, **(H)** *Dist_sf_*, **(I)** SLA, **(J)** *S_if_*, **(K)** *S_af_*, **(L)** *S_tr_*, **(M)** *S_sf_*.

**Supplementary Figure 5.**
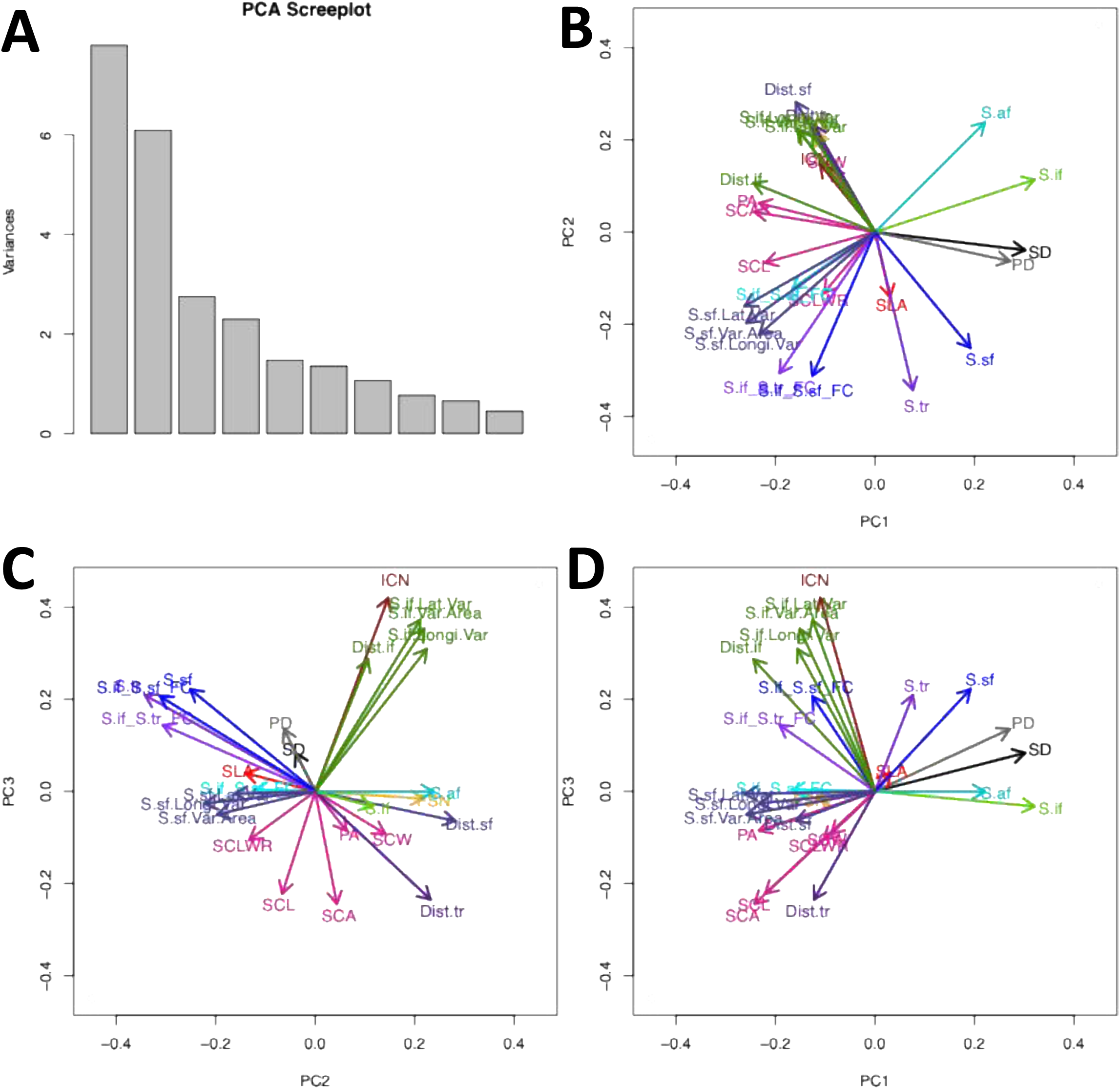
Principal components Analysis of the combined Xie, *et al*. and SPP traits. **(A)** Scree plot of the loadings for each PC dimension showing that the majority of the variance for these traits is found within PCs 1 and 2 with a large decrease by PC3. These loadings were then used to focus on which PCs were the most informative to assess and identify overlapping rotational loadings such as **(B)** the rotational loadings of PCs 1 and 2, **(C)** the rotational loadings of PCs 2 and 3, and **(D)** the rotational loadings of PCs 1 and 3.

**Supplementary Figure 6.**
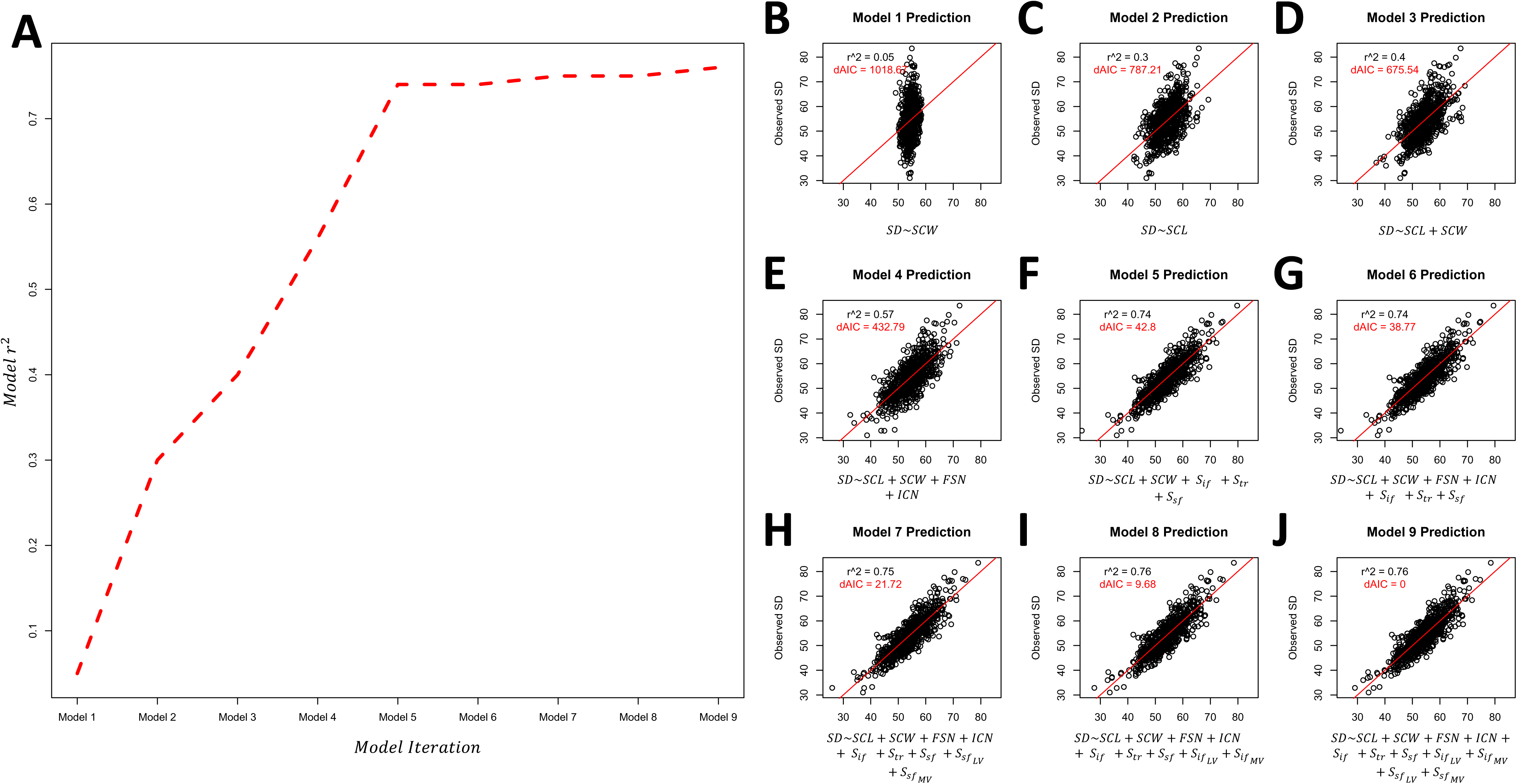
Linear model predictions of stomatal density for the 2016 field season inferred by using other associated traits as linear model coefficients with the r^2^ values shown in black for each model fit and corresponding AIC statistics denoted in red. **(B)** Model 1 estimates SD from stomatal widths, **(C)** Model 2 estimates SD from stomatal lengths, and **(D)** Model 3 estimates SD from the combined stomatal lengths and widths. Assuming the necessity of Model 3’s structure, **(E)** Model 4 further adds the stomatal file spacing number and asymmetric cell gap number, **(F)** Model 5 by contrast adds the In-File, Trench, and Side-File probabilities, and **(G)** Model 6 adds the combined influences of the stomatal file spacing number and asymmetric cell gap number and the In-File, Trench, and Side-File probabilities. Assuming the necessity of Model 6’s structure, **(H)** Model 7 adds the longitudinal and lateral variances of the side-file masks, by contrast **(I)** Model 8 adds the longitudinal and lateral variances of the side-file masks, and finally **(J)** Model 9 adds the combined influences of the longitudinal and lateral variances of both the in-file and side-file masks.

**Supplementary Figure 7.**
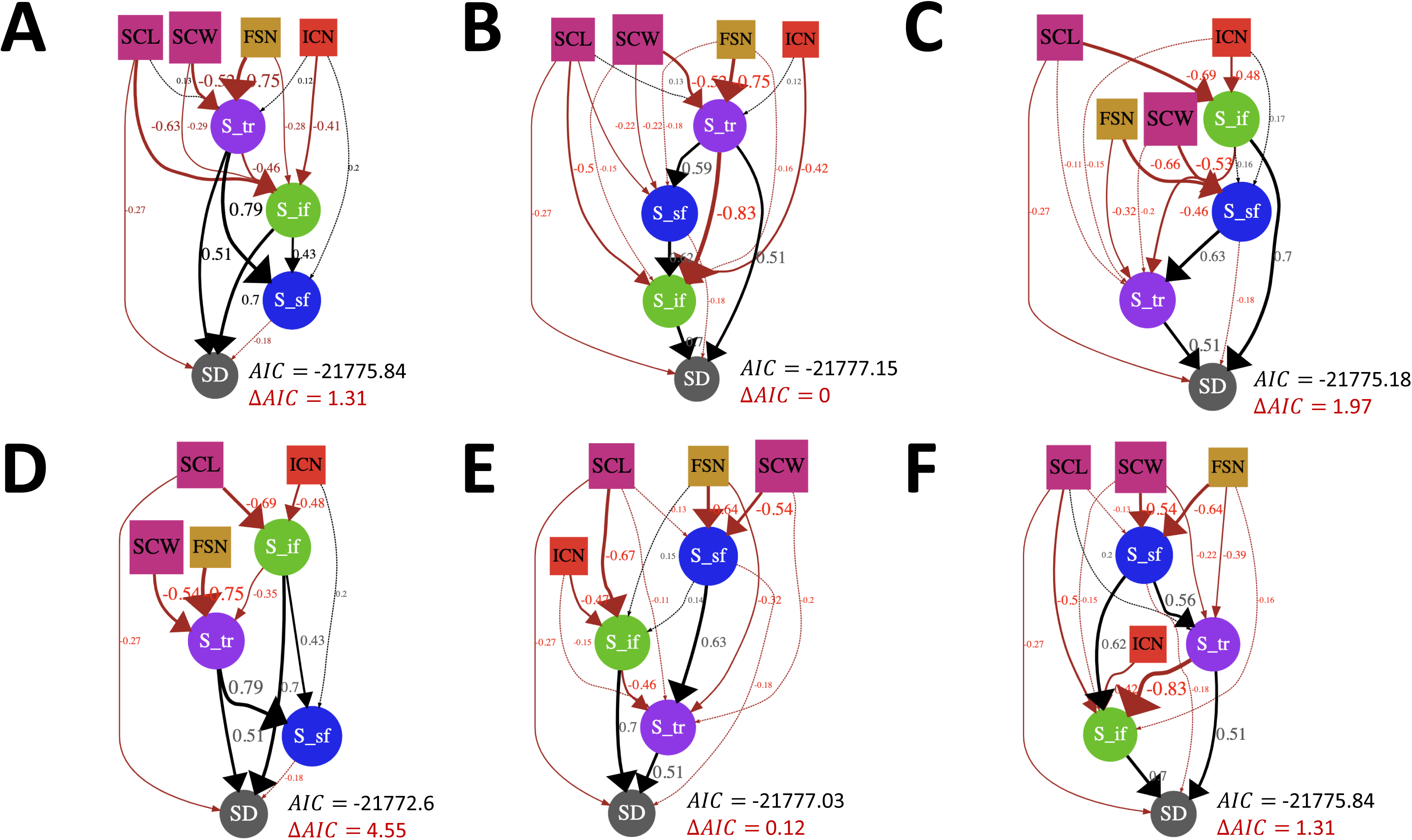
Alternative structural equation model designs with corresponding AIC statistics in which the optimal probability order was determined based on **(A)** trench → in-file → side-file, **(B)** trench → side-file → in-file, **(C)** in-file → **s**ide-file → trench, **(D)** in-file → trench → side-file, **(E)** side-file → in-file → trench, **(F)** side-file → trench → in-file causal hierarchies.

**Supplementary Figure 8.**
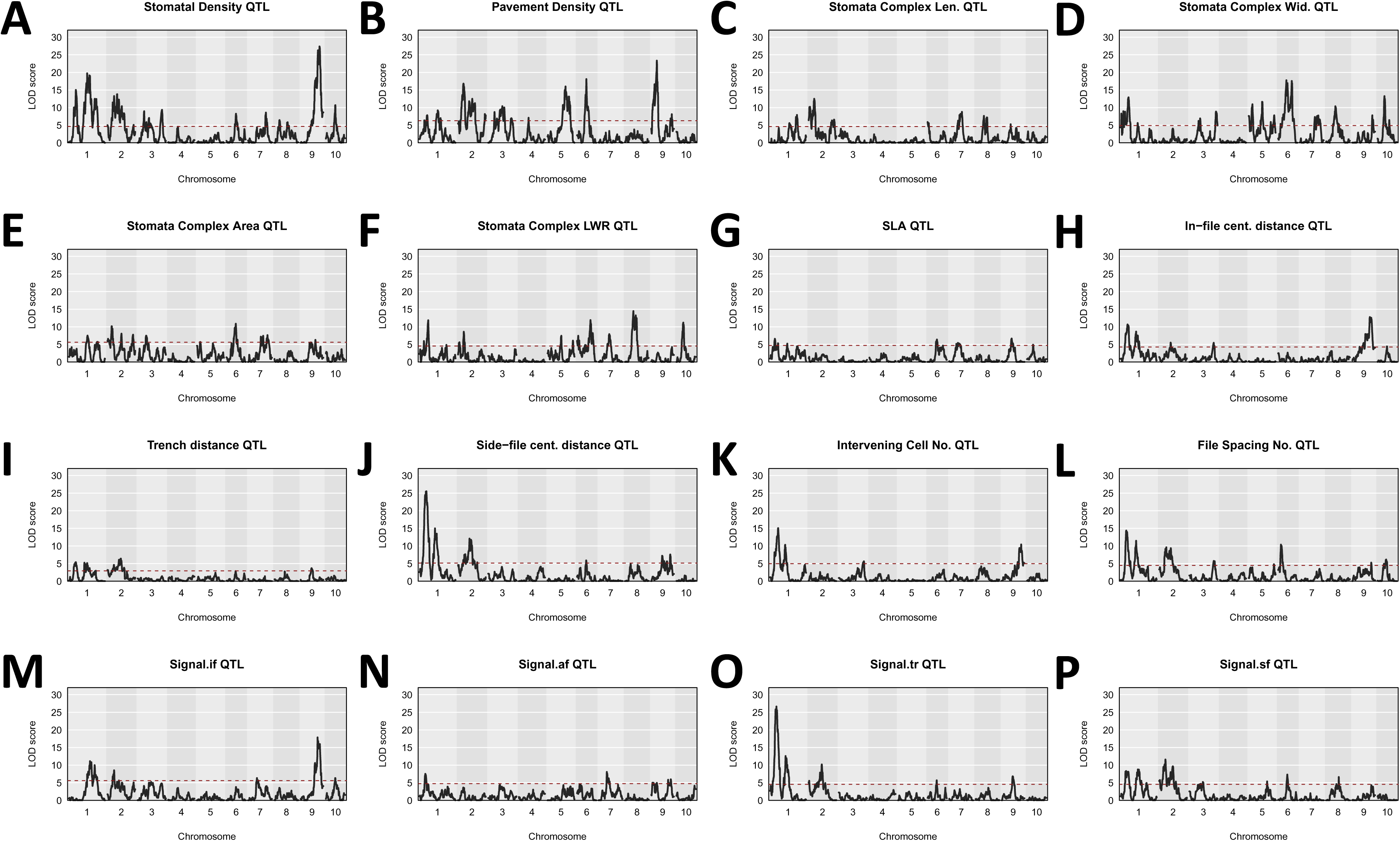
QTL LOD maps corresponding to the various traits screened for genetics associations across the maize population including **(A)** stomatal density, **(B)** pavement density, **(C)** stomatal complex length, **(D)** stomatal complex width, **(E)** stomatal complex area, **(F)** stomatal complex length: width ratio, **(G)** specific leaf area, **(H)** In-file centroid distance, **(I)** Trench distance, **(J)** side-file centroid distance, **(K)** intervening cell number, **(L)** file spacing number, **(M)** in-file probability, **(N)** diffuse-file probability, **(O)** trench probability, and **(P)** side-file probability. Alpha thresholds=0.0005 of are depicted as a dotted red line and were used to threshold large effect QTL for each QTL map.

**Supplementary Figure 9.**
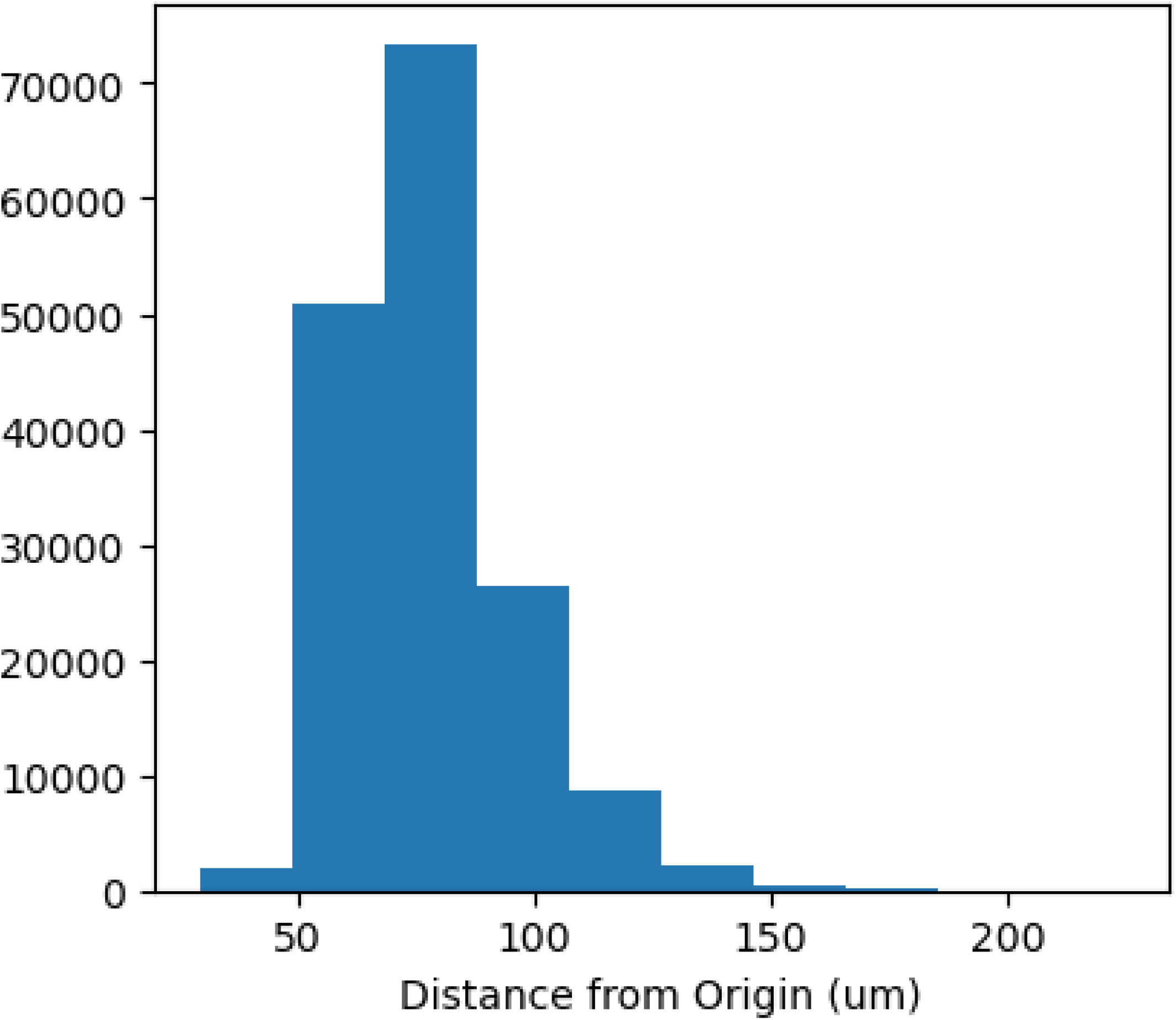
Examples from the maximum distances between the origin stomata and the 5^th^ ranked nearest neighbor used to define the x-y axis bounds for the SPPs.

## Supplementary Materials

The properties of the Manhattan distance formula always ensured a diamond-like pattern in both the NN and SPP heatmap data, however, a generalized null model compared against the B73 as a test case found this insufficient to capture the hotspots and gaps that were observed (**Supp. Fig. 2A-D**). The differences between these observed and expected kernel probabilities made these incongruences even more pronounced with two longitudinally biased “in-file hotspots” often with discernible tails radiating outwards (**Supp. Fig. 2E-G**). Two minor “side-file hotspots” also occur orthogonally to the in-file along the lateral plane with the “trench gap” being recognizable as a uninhabited space between these in-file and side-file regions. This trench wouldn’t be anticipated as an artifact of the Manhattan distance method (**Supp. Fig. 2F, G**). This data suggests that while the Manhattan method may help accentuate patterns like the in-file hotspots this model by itself is insufficient to characterize their shape as well as the gaps observed. This morphology is remarkably consistent across SPPs in our population with the in-file, trench, side-file regions always being present despite some variation in their distance to the origin or how focused they are in space.

## Notes

https://databank.illinois.edu/datasets/IDB-8275554

### Competing Interest Statement

The authors have declared no competing interest.

## References

1. Al-Salman Y, Cano FJ, Pan L, Koller F, Piñeiro J, Jordan D, Ghannoum O (2023). Anatomical drivers of stomatal conductance in sorghum lines with different leaf widths grown under different temperatures. Plant Cell Environ 46: 2142–58.

2. Ashraf, MA, Liu L, Facette MR (2023). A polarized nuclear position specifies the correct division plane during maize stomatal development. Plant Physiol 193: 125–39.

3. Aurenhammer F (1991). Voronoi diagrams – A survey of a fundamental geometric data structure. ACM Comput Surv 23(3): 345–405.

4. Balkunde R, Deneer A, Betchtel H, Zhang B, Herberth S, Pesch M, Jaegle B, Fleck C, Hülskamp M (2020). Identification of the trichome patterning core network using data from weak ttg1 alleles to constrain the model space. Cell Reports 33(108497): 1–14

5. Banerjee BP, Joshi S, Thoday-Kennedy E, Pasam RK, Tibbits J, Hayden M, Spangenberg G, Kant S (2020). High-throughput phenotyping using digital and hyperspectral imaging-derived biomarkers for genotypic nitrogen response. J Exp Bot 71(15): 4604–4615.

6. Baresch A, Crifò, CK (2019). Competition for epidermal space in the evolution of leaves with high physiological rates. New Phytologist 221: 628–639.

7. Broman KW, Gatti DM, Simecek P, Furlotte NA, Prins P, Sen S, Yandell BS, Churchill GA (2019). R/qtl2: Software for mapping quantitative trait loci with high-dimensional data and multiparent populations. Genet 211(2): 495–502.

8. Cao S, Xu D, Hanif M, Xia X, He Z (2020). Genetic architecture underpinning yield component traits in wheat. Theor Appl Genet 133: 1811–1823.

9. Casson S & Gray JE (2008). Influence of environmental factors on stomatal development. New Phytologist 178: 9–23.

10. Chen X, Liu L, Lee E, Han X, Rim Y, Chu H, Kim S, Sack F, Kim J (2009). The Arabidopsis Callose Synthase Gene GSL8 is required for cytokinesis and cell patterning. Plant Physiol 150: 105–113.

11. Conklin PA, Strable J, Li S, Scanlon MJ (2018). On the mechanisms and development of monocot and dicot leaves. New Phytol 221: 706–724.

12. Covarrubias-Pazaran G (2016). Genome-assisted prediction of quantitative traits using the R package *sommer*. PLoS One 11(6): e0156744.

13. Croxdale JL (2000). Stomatal patterning in Angiosperms. American J Bot 87(8): 1069–1080.

14. Cuthill JFH, Guttenberg N, Ledger S, Crowther R, Huertas B (2019). Deep learning on butterfly phenotypes tests evolution’s oldest mathematical model. Science Advances 5(eaaw4967): 1–11.

15. Delgado D, Sánchez-Bermejo E, de Marcos A, Martín-Jimenez C, Fenoll C, Alonso-Blanco C, Mena M (2019). A Genetic Dissection of Natural Variation for Stomatal Abundance Traits in Arabidopsis. Front Plant Sci 10(1392): 1–16.

16. de Boer HJ, Price CA, Wagner-Cremer F, Dekker SC, Franks PJ, Veneklaas EJ. (2016). Optimal allocation of leaf epidermal area for gas exchange. New Phytol 210: 1219–1228.

17. Devos KM & Gale MD (2000). Genome Relationships: the Grass Model in Current Research. Plant Cell 12: 637–646.

18. Doheny-Adams, T. Hunt L, Franks PJ, Beerling DJ, Gray JE (2012). Genetic manipulation of stomatal density influences stomatal size, plant growth and tolerance to restricted water supply across a growth carbon dioxide gradient. Phil Trans. Roy. Soc. London B 367(1588): 547–556.

19. Doll Y, Koga H, Tsukaya H. (2023). Experimental validation of the mechanism of stomatal development diversification. J Exp Bot 74(18): 5667–5681.

20. Doust AN & Diao X (2017). Genetics and Genomics of Setaria. Cham, Switzerland: Springer International Publishing.

21. Edwards, CE, Ewers BE, McClung CR, Lou P, Weinig C (2012). Quantitative variation in water-use efficiency across water regimes and its relationship with circadian, vegetative, reproductive, and leaf gas-exchange traits. Mol Plant 5(3): 653–668.

22. Esau K (1960). Anatomy of Seed Plants (2^nd^ Edition). New York, New York: John Wiley & Sons, Inc.

23. Facette MR, Park Y, Sutimantanapi D, Luo A, Cartwright HN, Yang B, Bennett EJ, Sylvester AW, Smith LG (2015). The SCAR/WAVE complex polarizes PAN receptors and promotes division asymmetry in maize. Nature Plants 1: 1–8.

24. Facette MR, Rasmussen CG, van Norman JM. (2019). A plane of choice: coordinating timing and orientation of cell division during plant development. Curr Opin Plant Biol 47: 47–55.

25. Failmezger H, Jaegle B, Schrader A, Hülskamp M, Tresch A (2013). Semi-automated 3D leaf reconstruction and analysis of trichome patterning from light microscope images. PLOS Computational Biology 9(4): e1003029.

26. Farquhar GD & Sharkey TD (1994). Photosynthesis and Carbon Assimilation. In: Boote KJ, Bennett JM, Sinclair TR, Paulsen GM (editors) Physiology and Determination of Crop Yield. ASA, CSSA, and SSSA, Madison, WI. 187-210.

27. Feldman MJ, Paul RE, Banan D, Barrett JF, Sebastian J, Yee M, Jiang H, Lipka AE, Brutnell TP, Dinneny JR, Leakey ADB, Baxter I (2017). Time dependent genetic analysis links field and controlled environment phenotypes in the model C4 grass Setaria. PLOS Genetics 13(6): e1006841.

28. Ferguson JN, Fernandes SB, Monier B, Miller ND, Allen D, Dmitrieva A, Schmuker P, Lozano R, Valluru R, Buckler ES, Gore MA, Brown PJ, Spalding EP, Leakey ADB. (2021). Machine learning-enabled phenotyping for GWAS and TWAS of WUE traits in 869 field-grown sorghum accessions. Plant Physiology 187(3): 1481–1500.

29. Ferguson JN, Schmucker P, Dmitrieva A, Quach T, Zhang T, Ge Z, Nersesian N, Sato SJ, Clemente TE, Leakey ADB (2024). Reducing stomatal density by expression of a synthetic epidermal patterning factor increases leaf intrinsic water use efficiency and reduces plant water use in a C4 crop. J Exp Bot 75(21): 6823–36.

30. Fernandes SB & Lipka AE (2020). simplePHENOTYPES: Simulation of pleiotropic, linked, and epistatic phenotypes. BMC Bioinform 21(491): 1–10.

31. Freeling, M (1992). A conceptual framework for maize leaf development. Dev Biol 153: 44–58. doi: 10.1016/0012-1606(92)90090-4

32. Furbank RT & Tester M (2011). Phenomics – technologies to relieve the phenotyping bottleneck. Trends Plant Sci 16(12): 635–644.

33. Gailing O, Langenfeld-Heyser R, Polle A, Finkeldey R (2008). Quantitative trait loci affecting stomatal density and growth in Quercus robus progeny: implications for the adaptation to changing environments. Glob Change Biol 14: 1934–1946.

34. Garvin DF, Gu Y, Hasterok R, Hazen SP, Jenkins G, Mockler TC, Mur LAJ, Vogel JP (2008). Development of genetic and genomic research resources for Brachypodium distachyon, a new model system for grass crop research. Crop Sci 48(S1): 69–84.

35. Grotzinger AD, Rhemtulla M, de Vlaming R, Ritchie SJ, Mallard TT, Hill WD, Ip HF, Marioni RE, McIntosh AM, Deary IJ, Koellinger PD, Harden KP, Nivard MG, Tucker-Drob EM (2019). Genomic structural equation modeling provides insights into the multivariate genetic architecture of complex traits. Nat Hum Behav 3: 513–25.

36. Guo X, Park CH, Wang Z, Nickels BE, Dong J (2021). A spatiotemporal molecular switch governs plant asymmetric cell division. Nature Plants 7: 667–680.

37. Guseman JM, Lee JS, Bogenshutz NL, Peterson KM, Virata RE, Xie B, Kanaoka MM, Hong Z, Torii KU (2010). Dyregulation of cell-to-cell connectivity and stomatal patterning by loss-of-function mutation in Arabidopsis CHORUS (Glucan Synthase-like 8). Development 137: 1731–41.

38. Hansen TF, Pélabon C, Houle D (2011). Heritability is not Evolvability. Evol Biol 38: 258–277.

39. Holbrook NM & Zweinicki MA (2005). Vascular Transport in Plants. Burlington, Massachusetts: Elsevier Academic Press.

40. Hughes TE & Langdale JA (2022). SCARECROW is deployed in distinct contexts during rice and maize leaf development. Development 149: 1–11.

41. John GP, Scoffoni C, Buckley TN, Villar R, Poorter H, Sack L (2017). The anatomical and compositional basis of leaf mass per area. Ecol Lett 20: 412–25.

42. Kaliman S, Jayachandran C, Rehfeldt F, Smith A (2016). Limits of applicability of the Voronoi tessellation determined by the centers of cell nuclei to epithelium morphology. Front Physiol 7(551): 1–14.

43. Kamiya N, Itoh J, Morikami A, Nagato Y, Matsuoka M (2003). The SCARECROW gene’s role in asymmetric cell divisions in rice plants. Plant Journal 36: 45–54.

44. Keller JM, Gray MR, Givens jr. JA (1985). A Fuzzy K-nearest neighbor algorithm. IEEE SMCS 15(4): 580–585.

45. Kellogg EA (2013). C4 photosynthesis. Current Biology 23(14): 594–599

46. Kovács AL. (1990). Hierarchical Processes in Biological Systems. Mathl Comp Modelling 14: 674–679.

47. Kumar D & Kellogg EA (2018). Getting closer: vein density in C4 leaves. New Phytologist 221: 1260–1267.

48. Lawson T & McElwain JC (2016). Evolutionary trade-offs in stomatal spacing. New Phytologist 210(4): 1149–1151.

49. Lawson & Leakey (2024). Stomata: custodians of leaf gaseous exchange. J Exp Bot 75(21): 6677–82.

50. Lau OS & Bergmann DC (2012). Stomatal development: a plant’s perspective on cell polarity, cell fate transitions and intercellular communication. Development 139: 3683–3692.

51. Leakey ADB, Ferguson JN, Pignon CP, Wu A, Jin Z, Hammer GL, Lobell DB (2019). Water use efficiency as a constraint and target for improving the resilience and productivity of C3 and C4 crops. Ann Rev Plant Biol 70: 781–808.

52. Lefcheck JS (2015). PiecewiseSEM: Piecewise Structural Equation Modelling in R for ecology, evolution, and systematics. Methods Ecol Evol 7(5): 573–79.

53. Li R, Tsaih S, Shockley K, Stylianou IM, Wergedal J, Paigen B, Churchill GA (2006). Structural model analysis of multiple quantitative traits. PLOS Genetics 2(7): 1046–1057.

54. Li J, Glover JD, Zhang H, Peng M, Tan J, Mallick CB, Hou D, Yang Y, Wu S, Liu Y, Peng Q, Zheng SC, Crosse EI, Medvinsky A, Anderson RA, Brown H, Yuan Z, Zhou S, Xu Y, Kemp JP, Ho YYW, Loesch DZ, Wang L, Li Y, Tang S, Wu X, Walters RG, Lin K, Meng R, Lv J, Chernus JM, Neiswanger K, Feingold E, Evans DM (2022). Limb development genes underlie variation in human fingerprint patterns. Cell 185: 95–112.

55. Lunn D, Kannan B, Germon A, Leverett A, Clemente TE, Altpeter F, Leakey ADB (2024). Greater aperture counteracts effects of reduced stomatal density on water use efficiency: a case study on sugarcane and meta-analysis. J Exp Bot **erae**271: 1–13.

56. MacAlister CA, Ohashi-Ito K, Bergmann DC. (2007). Transcription factor control of asymmetric cell divisions that establish the stomatal lineage. Nature 445: 537–540.

57. Maimaitijiang M, Sagan V, Erkbol H, Adrian J, Newcomb M, LeBauer D, Pauli D, Shakoor N, Mockler TC (2020). UAV-based Sorghum growth monitoring: a comparative analysis of LiDAR and photogrammetry. ISPRS Annals **V****-****3-**2020 489–496.

58. McKown KH & Bergmann DC (2020). Stomatal development in the grasses: lessons from models and crops (and crop models). New Phytologist 227: 1636–1648.

59. Mi X, Eskridge K, Wang D, Baenziger S, Campbell T, Gill KS, Dweikat I, Bovaird J (2010). Regression-based multi-trait QTL mapping using structural equation modeling. (2010). Stat Appl Genet Mol Biol 9(1): 1-23.

60. Mullet J, Morishige D, McCormick R, Truong S, Hilley J, McKinley B, Anderson R, Olson SN, Rooney W (2014). Energy Sorghum – a genetic model for the design of C4 grass bioenergy crops. J Exp Bot 65(13): 3479–3489.

61. Niklas KJ (1994). Plant allometry: Scaling form and process. Chicago, Illinois: University of Chicago Press.

62. Nelissen H, Rymen B, Jikumaru Y, Demuynck K, van Lijsebettens M, Kamiya Y, Inzé, Beemster GTS (2012). A local maximum in gibberellin levels regulates maize leaf growth by spatial control of cell division. Curr Biol 22: 1183–1187.

63. Nunes TDG, Zhang D, Raissig MT (2020). Form, development and function of grass stomata. Plant Journal 101: 780–799.

64. Ottaviano E & Camussi A (1981). Phenotypic and genetic relationships between yield components in maize. Euphytica 30: 601–609.

65. Raissig MT, Abrash E, Bettadapur A, Vogel JP, Bergmann DC (2016). Grasses use an alternatively wired bHLH transcription factor network to establish stomatal identity. PNAS 113(29): 8326–8331.

66. Raissig MT, Matos JL, Gil MXA, Kornfeld A, Bettadapur A, Abrash E, Allison HR, Badgley G, Vogel JP, Berry JA, Bergmann DC (2017). Mobile MUTE specifies subsidiary cells to build physiologically improved grass stomata. Science 355: 1215–1218.

67. Rajurkar AB, McCoy SM, Ruhter J, Mulcrone J, Freyfogle L, Leakey ADB (2022). Installation and imaging of thousands of minirhizotrons to phenotype root systems of field-grown plants. Plant Methods 18(39): 1–12.

68. Reynolds BE (1980). Taxicab Geometry. Pi Mu Epsilon J 7(2): 77–88.

69. Richardson AE, Cheng J, Johnston R, Kennaway R, Conlon BR, Rebocho AB, Kong H, Scanlon MJ, Hake S, Coen E (2021). Evolution of the grass leaf by primordium extension and petiole lamina remodeling. Science 374(6573): 1377–1380.

70. Riska B (1989). Composite traits, selection response, and evolution. Evolution 43(6): 1172–1191.

71. Sack L & Buckley TN (2016). The developmental basis of stomatal density and flux. Plant Physiol 171: 2358–2363.

72. Sage RF & Monson RK (1999). C4 plant biology. Academic Press, New York, New York. 133-172.

73. Saibo NJM, Vriezen WH, Beemster GTS, van der Straeten D (2006). Plant J 33: 989–1000.

74. Sedelnikova OV, Hughes TE, Langdale JA (2018). Understanding the genetic basis of Kranz anatomy with a view to engineering C3 crops. Annu Review Genet 52: 249–70.

75. Schuler ML, Sedelnikova OV, Walker BJ, Westhoff P, Langdale JA (2018). SHORTROOT-mediated increase in stomatal density has no impact on photosynthetic efficiency. Plant Physiology 176: 757–772.

76. Shipley B (2000). Cause and correlation in Biology: A User’s Guide to Path Analysis, Structural Equations and Causal Inference. Cambridge, England: Cambridge Univ. Press.

77. Sigmon B & Vollbrecht E (2010). Evidence of selection at the *ramosa1* locus during maize domestication. Mol Ecol 19: 1296–1311.

78. Slewinski TL, Anderson AA, Zhang C, Turgeon R (2013). Scarecrow plays a role in establishing Kranz Anatomy in Maize leaves. PCP 53(12): 2030–2037.

79. Slewinski TL, Anderson AA, Price S, Withee JR, Gallagher K, Turgeon R (2014). Short-root1 plays a role in the development of vascular tissue and Kranz anatomy in Maize leaves. Molecular Plant 7: 1388–1392.

80. Snowder GD & Fogarty NM. (2009). Composite trait selection to improve reproduction and ewe productivity: a review. Anim Prod Sci 49: 9–16.

81. Spiegelhalder RP & Raissig MT (2021). Morphology made for movement: formation of diverse stomatal guard cells. Curr Opin Plant Biol 63: 1–10.

82. Stebbins GL & Shah SS (1960). Developmental studies of cell differentiation in the epidermis of monocotyledons. Developmental Biology 2: 477–500.

83. Stotz GC, Salgado-Luarte C, Escobedo VM, Valladares F, Gianoli E (2022). Phenotypic plasticity and the leaf economics spectrum: plasticity is positively associated with specific leaf area. Oikos **e**09342: 1–7.

84. Stuber CW, Edwards MD, Wendel JF (1987). Molecular marker-facilitated investigations of quantitative trait loci in Maize. II. Factors influencing yield and it’s component traits. CBGC 27: 639–648.

85. Tan GD, Chaudhuri U, Varela S, Ahuja N, Leakey ADB (2024). Machine learning-enabled computer vision for plant phenotyping: a primer on AI/ML and a case study on stomatal patterning. J Exp Bot 75(21): 6683–6703.

86. Templeton AR (2006). Population Genetics and Microevolutionary Theory. Hoboken, New Jersey: John Wiley and Sons Inc.

87. Torii KU (2021). Stomatal development in the context of epidermal tissues. Ann Bot 128: 137–148.

88. Trewavas A. (2006). A Brief History of Systems Biology. Plant Cell 18: 2420–2430.

89. Vőfély RV, Gallagher J, Pisano GD, Bartlett M, Braybrook SA (2018). Of puzzles and pavements: a quantitative exploration of leaf epidermal cell shape. New Phytol 221: 540–552.

90. Wagner GP, Laubichler MD, Bagheri-Chailian H (1998). Genetic theory of epistatic effects. Genetica 102/103: 569–580.

91. Wei T & Simko V (2024). R package ‘corrplot’: visualization of a correlation matrix (version 0.95). url: https://github.com/taiyun/corrplot

92. Wang J, Lin Z, Zhang X, Liu H, Zhou L, Zhong S, Li Y, Zhu C, Lin Z (2019). Krn1, a major quantitative locus for kernel row number in maize. New Phytol 223(3): 1634–1646.

93. Wright, S (1921). The relative importance of heredity and environment in determining the piebald pattern of guinea-pigs. Genetics 6: 320–332.

94. Xie J, Fernandes SB, Mayfield-Jones D, Erice G, Choi M, Lipka AE, Leakey ADB (2021). Optical tomography and machine learning to rapidly phenotype stomatal patterning traits for maize QTL mapping. Plant Physiology 187(3): 1462-1480

95. Yang W, Guo Z, Huang C, Duan L, Chen G, Jiang N, Fang W, Feng H, Xie W, Lian X, Wang G, Luo Q, Zhang Q, Liu Q, Xiong L. (2014). Combining high-throughput phenotyping and genome-wide association studies to reveal natural genetic variation in rice. Nat Comm 5(5087): 1–9.

96. Yoo CY, Pence HE, Jin JB, Miura K, Gosney MJ, Hasegawa PM, Mickelbart MV (2010). The Arabidopsis GTL1 transcription factor regulates water use efficiency and drought tolerance by modulating stomatal density via transrepression of SDD1. Plant Cell 22: 4128–41.

97. Zeng SM, Lo EKW, Hazelton BJ, Morales MF, Torii KU (2020). Effective range of non-cell autonomous activator and inhibitor peptides specifying plant stomatal patterning. Development 147: 1–13.

98. Zhang JY, He SB, Li L, Yang HQ (2014). Auxin inhibits stomatal development through MONOPTEROS repression of a mobile peptide gene STOMAGEN in mesophyll. PNAS 111: E3015-E3023.

99. Zhou W, Yin J, Zhou Y, Li Y, He H, Yang Y, Wang X, Lian X, Dong X, Ma Z, Chen L, Hou S (2025). DSD1/ZmICEb regulates stomatal development and drought tolerance in maize. J Integr Plant Biol 0(0): 1–14 (Early access)

